# Profilin and formin constitute a pacemaker system for robust actin filament growth

**DOI:** 10.1101/736272

**Authors:** Johanna Funk, Felipe Merino, Larisa Venkova, Pablo Vargas, Stefan Raunser, Matthieu Piel, Peter Bieling

## Abstract

The actin cytoskeleton drives many essential biological processes, from cell morphogenesis to motility. Assembly of functional actin networks requires control over the speed at which actin filaments grow. How this can be achieved at the high and variable levels of soluble actin subunits found in cells is unclear. Here we reconstitute assembly of mammalian, non-muscle actin filaments from physiological concentrations of profilin-actin. We discover that under these conditions, filament growth is limited by profilin dissociating from the filament end and the speed of elongation becomes insensitive to the concentration of soluble subunits. Profilin release can be directly promoted by formin actin polymerases even at saturating profilin-actin concentrations. We demonstrate that mammalian cells indeed operate at the limit to actin filament growth imposed by profilin and formins. Our results reveal how synergy between profilin and formins generates robust filament growth rates that are resilient to changes in the soluble subunit concentration.

## Introduction

Eukaryotic cells move, change their shape and organize their interior through dynamic actin networks. Actin assembly requires nucleation of filaments, which elongate by the addition of subunits to filament ends. To move and quickly adapt their shape, most eukaryotic cells sustain vast amounts (>50uM) of polymerizable subunits, which requires the monomer-binding protein profilin (Koestler et al., 2009; Pollard et al., 2000; Raz-Ben Aroush et al., 2017; Skruber et al., 2018). Profilin shields the barbed end side of actin monomers to suppress spontaneous nucleation (Schutt et al., 1993). This allows profilin-actin complexes to exist at high concentration *in vivo*, unlike free actin monomers. Profilin-actin is therefore considered the physiological substrate of filament growth (Kaiser et al., 1999; Pantaloni and Carlier, 1993; Pollard et al., 2000) which occurs when profilin-actin complexes bind to exposed filament barbed ends (Gutsche-Perelroizen et al., 1999; Kinosian et al., 2002; Pollard and Cooper, 1984; Pring et al., 1992). The speed of filament elongation over a limited concentration range of profilin-actin fits a linear model for a binding-controlled reaction (Blanchoin and Pollard, 2002; Oosawa and Asakura, 1975). This has led to the idea that the concentration of soluble subunits is the central parameter that controls the speed of actin growth (Blanchoin et al., 2014; Carlier and Shekhar, 2017; Pollard et al., 2000). However, actin elongation has only been studied at low, non-physiological levels of soluble subunits until now.

The concentration of profilin-actin is thought to pace not only spontaneous, but also catalyzed actin growth by actin polymerases such as formins (Paul and Pollard, 2009). These modular proteins bind the filament barbed end via their FH2 domain and recruit many profilin-actin complexes through flexible FH1 domains (Fig. 1A). Polymerase activity is thought to arise from formins ability to increase the binding frequency of profilin-actin (Courtemanche, 2018; Paul and Pollard, 2009; Vavylonis et al., 2006). Whether, however, filament growth *in vivo* is controlled at the level of binding is unknown. Consequently, we do not fully understand how formins function as actin polymerases in cells.

**Fig. 1:**
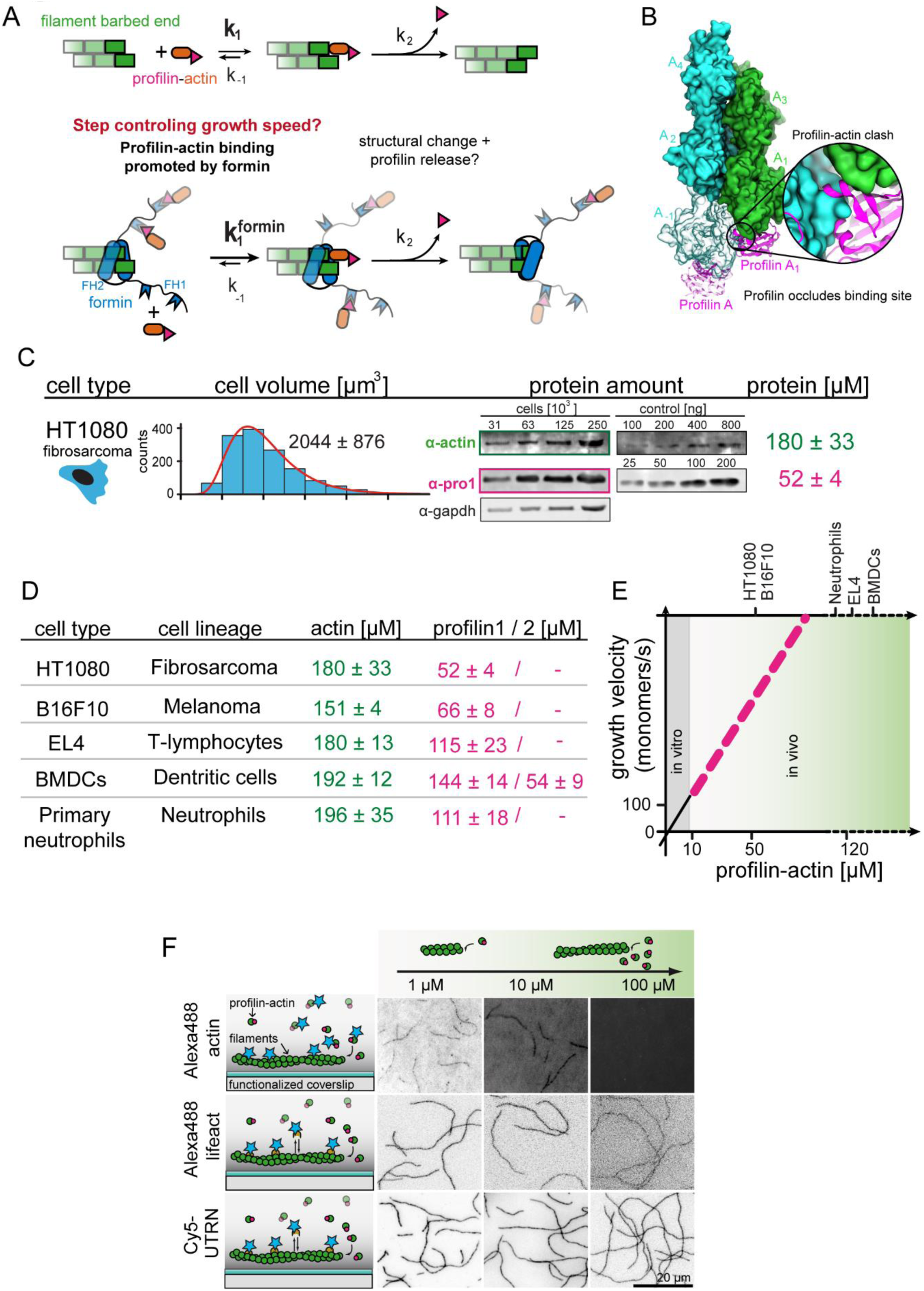
Filament assembly at physiological profilin-actin concentrations. (A) Scheme of barbed end elongation from profilin-actin alone (top) or with formins (bottom). (B) Structural model of profilin at filament barbed ends (Methods). The incoming profilin-actin complex is transparent. Actin is shown as green surface while profilin as magenta ribbons. Inset highlights the clash between the incoming actin monomer and profilin. (C) Profilin-actin measurements in HT1080 cells. Left to right: single cell volume histogram, western blots of actin, profilin1 (left: cell titration, right: standard curve of recombinant proteins), values are mean (N=3) and SD, Methods. (D) Table of total concentrations of actin and profilin-1/2 in various mammalian cell types (Fig. S1). (E) Scheme of a linearly substrate-dependent actin elongation rate. Top axis: Profilin-actin amounts for various cell types as indicated. (F) Scheme (left) and TIRFM images (right) of elongating filaments at indicated profilin-actin concentrations visualized with top-Alexa488-labeled monomers (20% labeled), middle - 10 nM Alexa488-lifeact, bottom – 10 nM Cy5-UTRN_261_.

The model of linear concentration-dependent scaling of actin growth creates a conundrum because of two reasons: i) Filament growth from profilin-actin complexes cannot occur in a single binding step, but requires additional reactions whose rate should not depend on the free subunit concentration (Fig.1A). Binding of profilin-actin to the actin filament barbed end occludes the binding site for new subunits and profilin needs to be released for elongation to continue (Fig. 1B) (Courtemanche and Pollard, 2013; Pernier et al., 2016; Pollard and Cooper, 1984). How rapidly profilin release occurs and whether it affects filament growth is presently unclear (Blanchoin and Pollard, 2002; Gutsche-Perelroizen et al., 1999; Romero et al., 2004). ii) Generally, soluble actin concentrations vary significantly across species, cell types (Koestler et al., 2009; Pollard et al., 2000; Raz-Ben Aroush et al., 2017) and likely even within a single cell (Skruber et al., 2018). If elongation rates scale linearly with profilin-actin concentrations, then actin filaments must grow at widely different speeds *in vivo*. Actin polymerases like formins should dramatically amplify such variations. This poses a challenge to the construction of functional actin networks whose architecture depends on the filament elongation speed. We presently do not understand how cells control the rate of filament growth when facing variable and fluctuating profilin-actin levels. Here we uncover a mechanism that establishes robust, but tunable growth rates that are buffered against changes in the free subunit concentration.

## Results

### Actin filament growth at physiological profilin-actin concentrations

To reconstitute actin assembly at cell-like conditions, we first determined the concentration of actin and the two most abundant profilin isoforms (−1 and −2) (Mouneimne et al., 2012) in mammalian cells through volume measurements (Cadart et al., 2017) and western blots (Fig. 1C-D & S1A, Methods). We studied mesenchymal (HT1080), epithelial (B16F10) or immune cells (T-cells, dendritic cells and neutrophils), the latter because of their rapid motility. Consistent with earlier estimates (Pollard et al., 2000; Witke et al., 2001) profilin and actin were highly expressed (Fig. 1D, S1A). Profilin-1 was the dominant isoform, whereas profilin-2 was not present at substantial levels in most cells (Fig. S1A). Profilin levels were especially high in immune cells, in line with their dynamic actin cytoskeleton. Actin always exceeded the profilin concentration as expected, since actin forms filaments and binds monomer-binding proteins other than profilin. Because profilin binds mammalian cytoplasmic actin much more tightly than other abundant monomer-binding proteins like thymosin-β_4_ (see below), the actin pool is likely sufficiently large for profilin to be nearly completely bound to monomers *in vivo* (Kaiser et al., 1999). We thus estimated the profilin-actin concentration around 50 - 200 μM, depending on mammalian cell type (Fig. 1D-E).

To study actin elongation at these conditions, we first adapted methods (Hatano et al., 2018; Ohki et al., 2009) to purify mammalian cytoplasmic actin (β-γ isoforms). Past studies relied mostly on muscle α-actin, the most divergent actin isoform. Its widespread use, combined with chemical labeling, created confusion concerning the role of profilin in the past (Blanchoin and Pollard, 2002; Courtemanche and Pollard, 2013; Kinosian et al., 2002, 2000; Pernier et al., 2016; Romero et al., 2007; Vavylonis et al., 2006). To study the authentic substrate of actin assembly in non-muscle cells, we purified either i) native bovine actin from thymus tissue or ii) recombinant human β-actin from insect cells (Fig. S1B, Methods). Using mass spectrometry, we detected β and γ actin in a roughly 1:1 ratio, but no α-actin in native actin (not shown). Mammalian cytoplasmic actin polymerized with rates comparable to actin from other organisms (Bieling et al., 2018; Pollard, 1986) (Fig. S1C).

We then studied binding of the most abundant monomer-binding proteins, profilin-1/-2 and thymosin-β_4_, to mammalian cytoplasmic actin (Fig. S1D-E). In general agreement with studies using non-muscle actin (Bieling et al., 2018; Kinosian et al., 2002; Vinson et al., 1998), thymosin-β_4_ bound weakly (*K_D_* ∼ 1.2 μM), whereas profilin bound exceptionally strongly (*K_D_* ∼ 18 nM) to actin monomers at near-physiological ionic strength. This allowed us to isolate profilin-actin complexes by size-exclusion chromatography (Fig. S1F) and to concentrate them (>500 μM) without triggering nucleation. We then turned to total internal-reflection fluorescence microscopy (TIRFM) assays to analyze elongation of surface-tethered actin filaments (Fig. 1F & 2A). Strong background prevented us from using fluorescent actin at high concentrations (Fig. 1F upper). Trace amounts (10 nM) of fluorescent filament-binding probes (UTRN_261_ or LifeAct), however, yielded sufficient contrast without altering assembly kinetics (Fig. 1F middle and lower, (Bieling et al., 2018)). To further minimize nucleation, we additionally added low amounts of either free profilin (<2 μM) or thymosin-β_4_ (<15 μM) at high profilin-actin concentrations (Methods). As expected, this did not impact filament elongation (Fig. S2A-B). These advances allowed us to, for the first time, study mammalian cytoplasmic actin growth over the entire physiological range of profilin-actin concentrations.

**Fig. 2:**
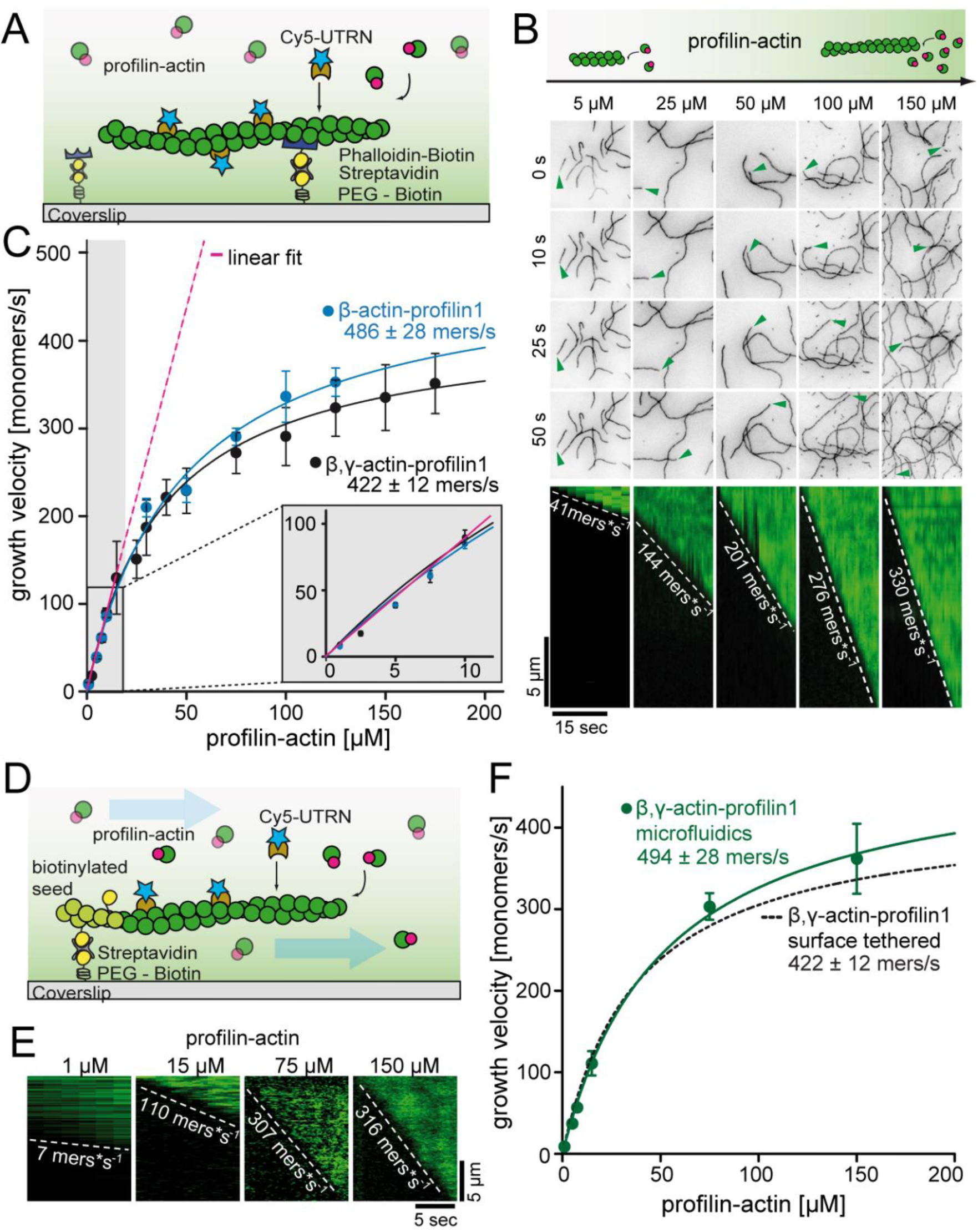
A kinetic limit to actin filament elongation from profilin-actin. (A) Scheme of TIRFM elongation assays of surface-attached filaments from profilin-actin on functionalized coverslips. (B) TIRFM time-lapse images (top) and kymographs (bottom) of filament elongation (green arrow follows a single barbed end) at indicated profilin-actin concentrations. (C) Barbed end growth velocities from TIRFM assays using different profilin1:actin complexes as indicated. Points are mean values [N ≥40 for each condition, error = SD]. Lines are hyperbolic fits. Inset: Regime of low concentrations fitted by a linear model (magenta, Fig. S2C-D). (D) Scheme of microfluidic experiments of seed-attached filaments under flow. (E) Kymographs of filaments at indicated profilin-actin concentrations in microfluidic experiments. (F) Barbed end growth velocities of filaments grown in microfluidic channels in TIRFM assays (green) compared to surface tethered filaments as quantified in ((C), black dashed line).

### Profilin dissociation kinetically limits filament elongation

As previously (Blanchoin and Pollard, 2002; Jégou et al., 2013), we observed a linear increase of the actin filament growth velocity at low profilin-actin concentrations (<10 μM, Fig. 2B-C & S2C-D). Strikingly, however, elongation rates deviated strongly from linearity at moderate (>20 μM) and nearly saturated at high (≥100 μM) concentrations to plateau at ∼400 monomers s^-1^ (Fig. 2B-C, Movie S1). Importantly, this maximal rate did not depend on surface tethering, the filament-binding probe or the specific cytoplasmic isoform of profilin or actin (Fig. S2C-E). We ruled out accumulation of free profilin as a reason for saturation, because filament growth was constant over time under all conditions (Fig. S2A-B). We observed saturation also in microfluidic assays with a constant influx of fresh profilin-actin for filaments that were only attached via short seeds (Fig. 2D-F & S2F). This demonstrates that filament elongation at physiological conditions is not controlled by binding of profilin-actin, but is kinetically limited by a reaction that proceeds with a rate of ∼400 s^-1^.

Structural models suggest that incorporation of profilin-actin transiently caps barbed ends, because profilin sterically hinders the binding of the next monomer (Fig. 1B, (Courtemanche and Pollard, 2013)). Profilin release is therefore required for continual elongation. Profilin binds much more weakly to filament barbed ends than to monomeric actin (Courtemanche and Pollard, 2013; Pernier et al., 2016). We confirmed that profilin dissociation from actin monomers (k_off_ = 0.77 s^-1^, Fig. S3B-D) is much slower than the maximal elongation rates we observe (∼400 s^-1^). This means that structural changes in the terminal actin protomer are required to trigger profilin release. We deduced that either of these subsequent reactions could become rate-limiting (Fig. 3A). Profilin dissociation specifically, should be affected by interactions between actin and profilin. To test this hypothesis, we introduced mutations in profilin-1 at the actin binding interface to either decrease (E82A, R88K) or increase (K125E+E129K) affinity (Fig. 3B-C, F & S3A, Methods). Single point mutants (E82A and R88K) caused a moderate reduction (∼ 1.5 and ∼ 4– fold, respectively), while mutation of two residues (K125E+E129K) showed an increase (∼ 5–fold) in monomer binding affinity (Fig. 3C). Importantly, these changes were caused by altered dissociation, but not association rate constants (Fig. S3B-C, 3F). More drastic changes were incompatible with elongation assays due to either accumulation of free actin (severely weakening mutants) or the complete inhibition of growth (ultra-tight binding mutants, Fig. S3E-F).

**Fig. 3:**
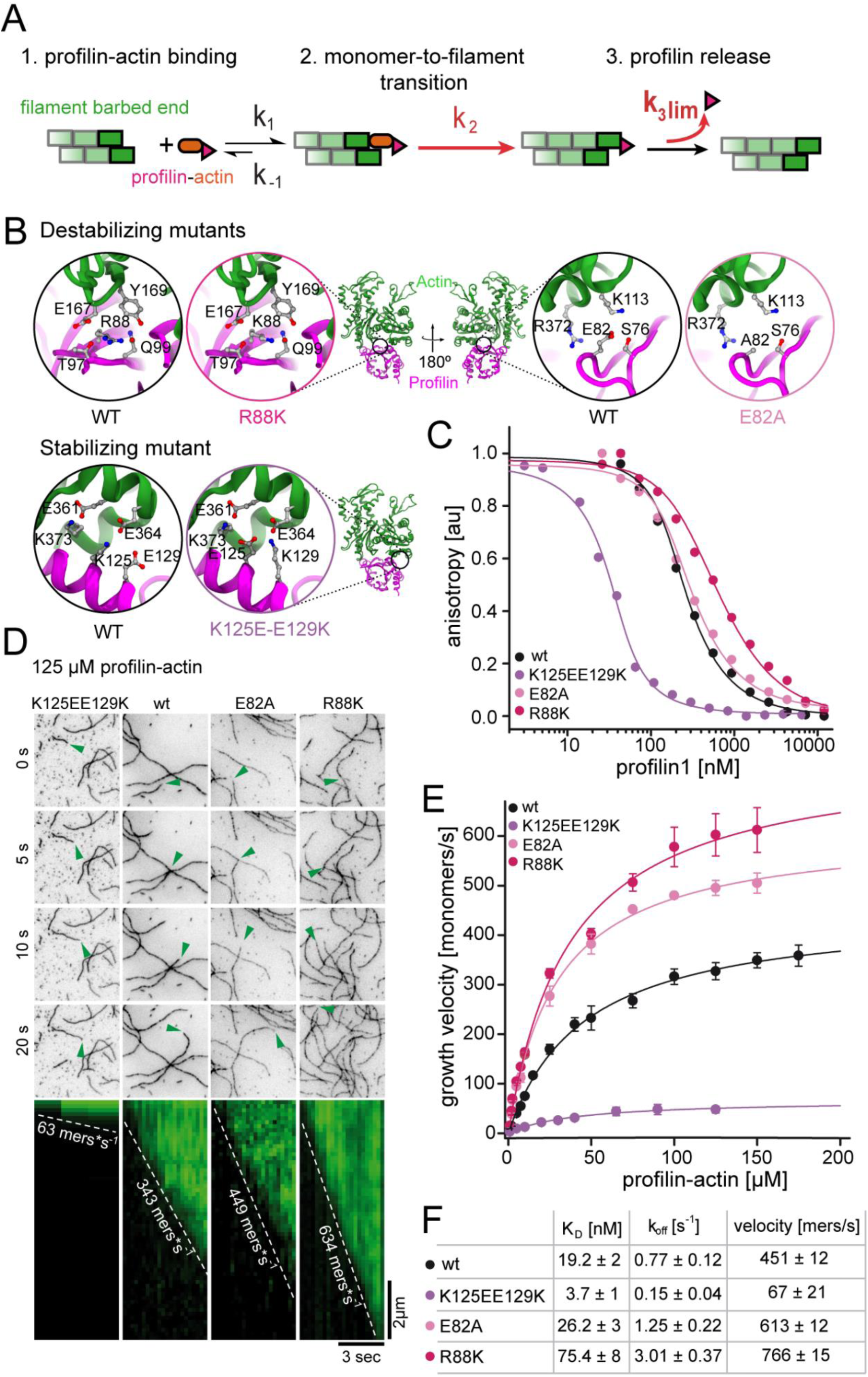
Profilin release kinetically limits filament elongation. (A) Scheme of barbed end elongation from profilin-actin alone indicating the potential limiting kinetic steps. (B) Structural models (Methods) of the actin interface of stabilizing and destabilizing profilin mutants. Ribbon diagrams highlight the mutation positions. Insets show changes in amino acid environments upon mutation. (C) Binding of profilin to actin monomers measured by fluorescence anisotropy competition assays. Fluorescence anisotropy of Atto488-WAVE1_WCA_ (4 nM) at increasing profilin1 (wt or mutants as indicated) concentrations in the presence of actin monomers (150 nM). Lines are fits to an analytical competition model (Methods). Points represents means (N ≥3) ± SD. (D) TIRFM time-lapse images (top) and kymographs (bottom) of filament elongation (green arrow follows a single barbed end) from mutant profilin1:actin complexes (125 μM total) as indicated. (E) Barbed end growth velocities measured from TIRFM assays using mutant profilin1:actin complexes as indicated. Points are mean values [N ≥40 for each concentration, error = SD]. Lines are hyperbolic fits. (F) Summary table of equilibrium dissociation constants (K_D_) and dissociation rate constants (k_off_, Fig. S3) of the interaction of profilin1 (wt or mutants as indicated) and actin monomers and the resulting maximal filament elongation velocities as measured by TIRFM.

We then tested the effect of these profilin mutations on filament growth. Strikingly, the maximal elongation rate scaled with the monomer dissociation rate of profilin. Weakly-binding profilins increased, whereas tight-binding profilin reduced the maximal filament growth rate (Fig. 3D-F, Movie S2). We draw two conclusions from these observations: i) The profilin mutations impact the dissociation of profilin from both soluble actin monomers and terminal actin subunits similarly. ii) The strength of the profilin-actin interaction modulates the rate-limiting step of elongation. This strongly suggests that profilin dissociation from the barbed end imposes a kinetic limit to actin filament elongation.

Some previous studies have linked profilin release from barbed ends to ATP hydrolysis within actin (Pernier et al., 2016; Romero et al., 2004). We therefore generated ATPase-deficient (AD) actin, by mutations of three residues within the catalytic core of actin (Q137A+D154A+H161A, Fig. 4A). These combined mutations did not abolish nucleotide binding, affect polymerization or reduce the affinity for profilin (Fig. 4B-E & S4A). Endpoint (Fig. 4B) and time-resolved ATPase assays (Fig. 4C) showed that this triple mutant did not hydrolyze ATP upon polymerization from profilin-actin. Importantly, we found that ATPase-deficient actin was able to elongate actin filaments with nearly the same rates as wildtype actin at saturating profilin-actin concentrations (Fig. 4D-E). This clearly demonstrates that profilin release from the barbed end does not require nucleotide hydrolysis in actin. More generally, this also suggest that ATP hydrolysis does not control actin assembly, but rather disassembly, most likely through actin binding proteins such as cofilin.

**Fig. 4:**
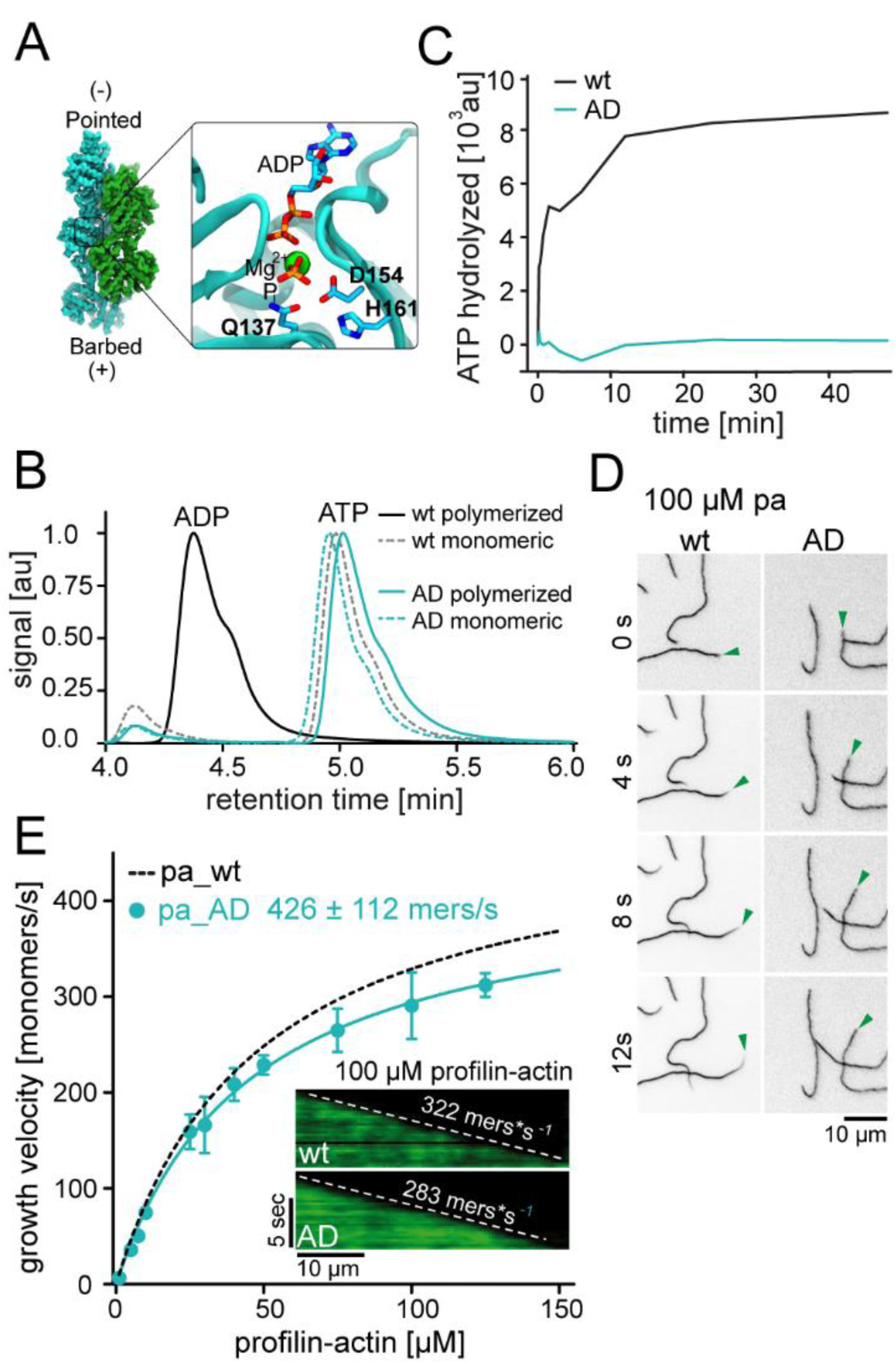
ATP hydrolysis is not required for profilin release from the barbed end. (A) Nucleotide-binding site of filamentous actin. Left: the overall structure of filamentous actin. Right: Inset of the active site (PDBID 6FHL), including the three amino acids involved in nucleotide hydrolysis, and the products of the reaction ADP and Pi. (B) End-point assays examining nucleotide content via HPLC after 1.5h of seeded polymerization from profilin-actin (either wt or AD). As a non-polymerized control, profilin-actin was stabilized via LatrunculinB before the experiment. (C) ATPase activity of wt and AD actin in seeded polymerization assays. The cleavage of γ-^32^P is monitored over time after mixing profilin1: actin complexes containing radioactive ATP with filaments in a 1:1 ratio (12 μM total)). (D) TIRF-M time-lapse images of filament barbed end elongation (green arrow follows a single barbed end) from either wt- or AD actin-containing profilin1-actin complexes (100 μM total). (E) Barbed end growth velocities of profilin1–actin (100 μM total, wt (black) or AD (cyan)) from TIRFM assays. Points are mean values [N ≥40 for each concentration, error = SD]. Lines are hyperbolic fits. Inset: Kymographs of filament growth.

### Formin actin polymerases promote profilin release through their FH2 domain

Actin elongation in cells can be facilitated by actin polymerases such as formins. These proteins are thought to increase the rate of binding between profilin-actin complexes and the barbed end they processively associate with (Paul and Pollard, 2009). Because such a mechanism can only accelerate growth when binding is limiting, we asked how formins affect actin assembly at saturating profilin-actin concentrations. We focused on Diaphanous-type formins because of their established polymerase function. We introduced constitutively active mDia1, containing profilin-actin-interacting FH1 and barbed end-binding FH2 domains, to TIRFM assays (Fig. 5A). We used formin concentrations sufficient to saturate filament barbed ends, as evident from their accelerated growth rate compared to control experiments (Methods). We verified that the measured velocities match the speed of formins observed at the single-molecule level (Movie S3). mDia1 strongly accelerated barbed-end growth at limiting profilin-actin concentrations (≤10 μM), as expected (Jégou et al., 2013; Kovar et al., 2006). Importantly, mDia1-mediated elongation still exhibited saturation at elevated profilin-actin levels, but converged to a much higher (4x-fold) maximal rate than observed for free ends (Fig. 5B-C & S5A). This demonstrates that formins can accelerate the rate-limiting reaction in filament elongation at saturating profilin-actin concentrations.

**Fig. 5:**
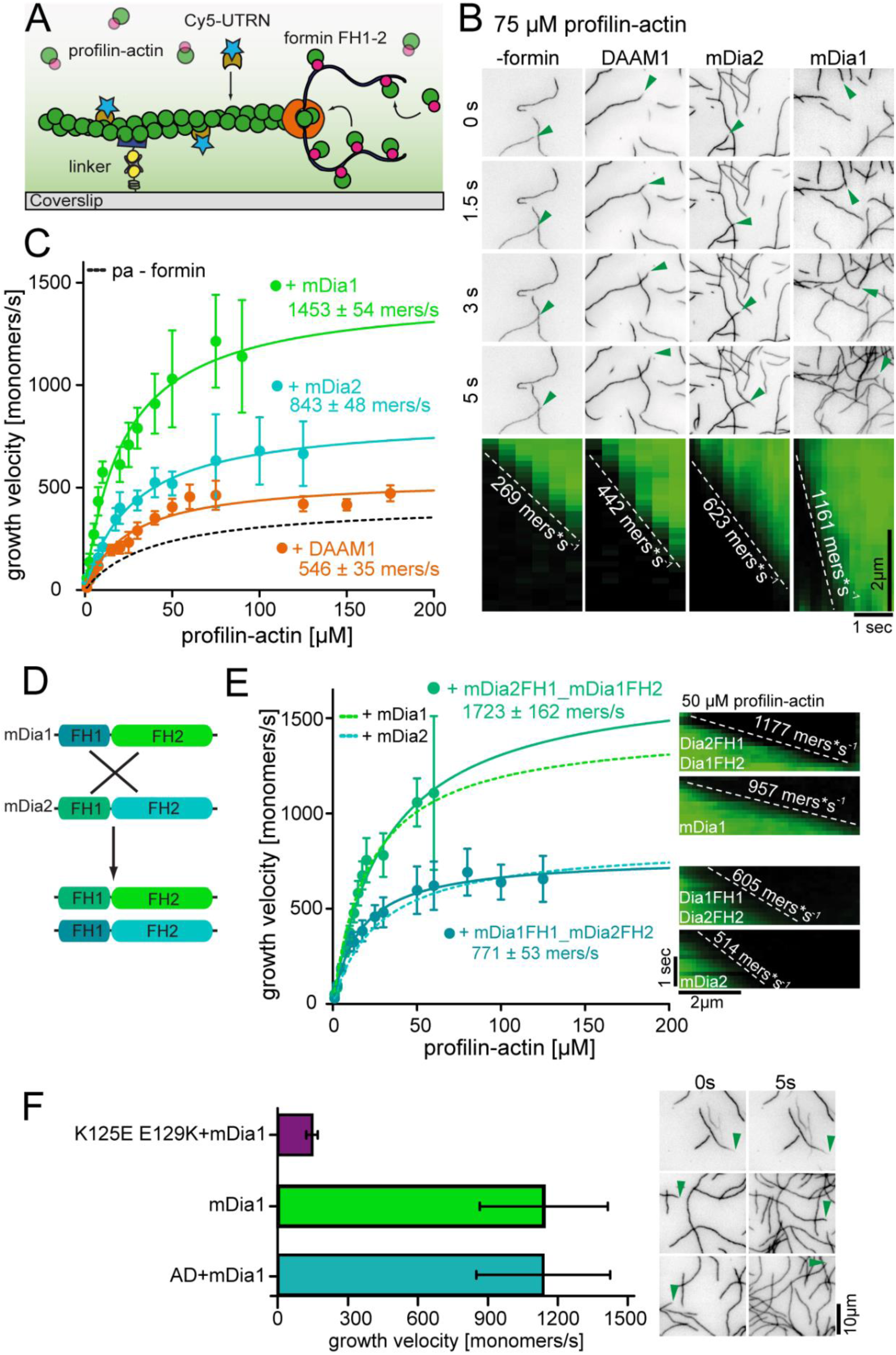
Formins accelerate filament elongation at saturating profilin-actin concentrations. (A) Scheme of TIRFM assays with formin catalyzing the elongation of a filament from profilin-actin on functionalized coverslips. (B) Top: TIRFM time-lapse images of formin-mediated actin elongation (green arrows follow a single barbed end) at 75 μM profilin-actin in the absence or in the presence of 15 nM formin constructs as indicated. Bottom: Kymographs of individual growing filaments as in the top panel. (C) Velocities of formin-catalyzed barbed end growth from TIRFM assays as in (B). Points are mean values [N ≥40 for each concentration, error = SD]. Lines are hyperbolic fits. (D) Scheme of the generation of mDia chimeras. (Methods). (E) Barbed end growth velocities of mDia chimeras (continuous lines) compared to wt mDia formins ((B), dashed lines) from TIRFM assays. Points are mean values [N ≥40 for each condition, error = SD]. Lines are hyperbolic fits. Right: Kymographs of growing filaments (± formins as indicated) at 50 μM profilin-actin. (F) Comparison of mDia1(15nM)-mediated filament growth in the 100 μM profilin-actin (either both wt proteins, tight binding profilin-1 (K125E-E129K) or ATPase-deficient actin (AD) as indicated). Left: Growth velocities. Right: TIRFM time-lapse images (green arrows follow a single barbed end).

To test whether this ability is shared among formins, we studied other diaphanous-(mDia2) and non-diaphanous (DAAM1) formins. Indeed, both mDia2 and DAAM1 accelerated filament elongation not only at limiting, but also saturating profilin-actin concentrations albeit less strongly than mDia1 (Fig. 5B-C, Movie S4). The relative rate enhancement of all formins decreased only slightly with substrate concentrations (Fig. S5A). Formins thus slightly broaden the regime over which actin growth is insensitive to the profilin-actin concentration (Fig. S5B).

Interestingly, even closely related formins such as mDia1 and 2 differ in their ability to accelerate the rate-limiting reaction of filament elongation. To understand the origin of this difference, we created chimeras of mDia1 and 2 by swapping their FH1 and FH2 domains (Fig. 5D). Both chimeras accelerated filament growth, but generated distinct maximal rates at saturating profilin-actin concentration (Fig. 5E). Interestingly, mDia2FH1-mDia1FH2 exhibited similar maximal rates as mDia1, whereas mDia1FH1-mDia2FH2 was comparable to mDia2 (Fig. 5E). This demonstrates that the barbed-end associated FH2 domain is responsible for setting the maximal rate of filament elongation.

Finally, to test which constraints limit formin-mediated growth, we elongated mDia1-associated actin filaments using profilin-actin complexes containing either ATPase-deficient actin or tight-binding profilin. ATPase-deficient actin grew with rates indistinguishable from wildtype actin, whereas tight profilin binding inhibited mDia1-mediated growth (Fig. 5F, S5C). This demonstrates that formin-mediated filament elongation at saturation is limited by profilin release from the barbed end and not nucleotide hydrolysis. These results uncover two distinct formin polymerase activities. Formins not only promote binding of profilin-actin complexes, but also directly accelerate profilin release from the barbed end via their FH2 domain. These activities are matched to provide a constant rate enhancement over a wide range of profilin-actin concentrations (Fig. S5A). Their combination allows formins to act as pacemakers, which elongate filaments with distinct rates that are buffered against changes in the profilin-actin concentration.

### Formin-mediated actin elongation is resilient to changes in profilin-actin levels

To critically test how our results relate to cellular actin growth, we sought to study actin filament elongation *in vivo*. Growth of individual actin filaments cannot be visualized in mammalian cells. Formin proteins, however, can be visualized as single molecules *in vivo* (Higashida et al., 2004). We thus established single-molecule TIRFM imaging of constitutively active, mNeonGreen-tagged formins within the cortex of either mammalian mesenchymal (HT1080) or T-lymphocyte (EL4) cells (Fig. 6A-C). We chose these cell types because of their >2-fold difference in profilin-actin levels (Fig. 1D). Because strong overexpression of active formins affects the soluble actin pool (Dimchev et al., 2017), we only analyzed cells with extremely low formin levels (Methods). Single formin molecules were visible as spots that translocated over μm distances with nearly constant velocity (Fig. 6B-C, Movie S5). Control experiments showed that formin movement was actin polymerization- and not myosin-driven (Movie S6). Remarkably, we observed that mDia1 and mDia2 moved with distinct speeds that were not only similar between the two cell types (Fig. 6D), but also strikingly close to their characteristic maximal *in vitro* velocity (1453 and 843 monomers/s for mDia1 and 2, respectively Fig. 5).

**Fig. 6:**
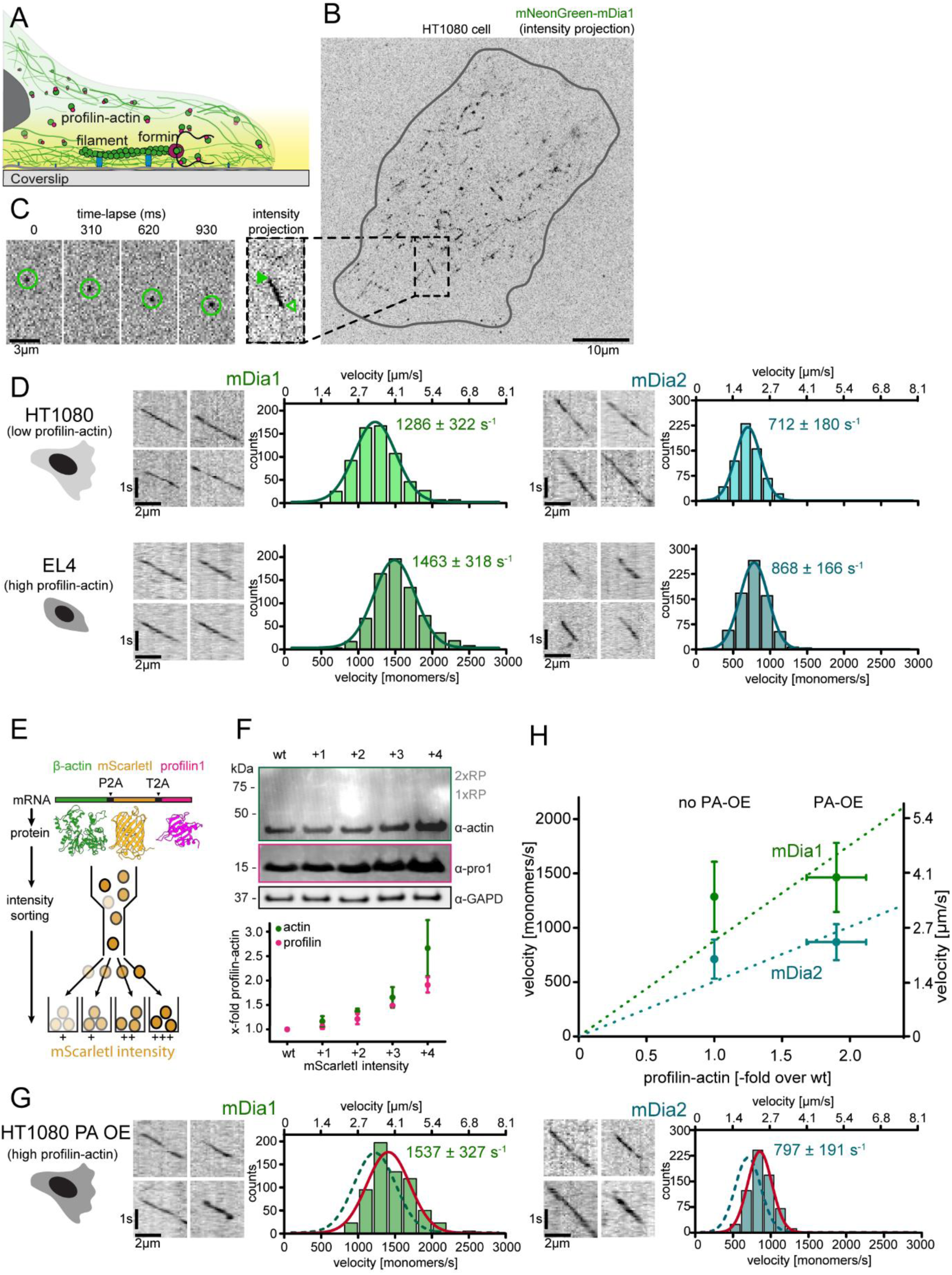
Formin single molecule imaging reveals buffered elongation rates in mammalian cells. (A) Scheme of TIRFM imaging of single formins in the actin cortex of cells. (B) Maximum intensity projection of a TIRFM time-lapse shows growth trajectories of single mNeonGreen-mDia1 molecules in the cortex of a HT1080 cell. Inset: Close-up of a single trajectory as in (C). (C) TIRFM time-lapse images (left) and intensity projection (right) of an individual mNeonGreen-mDia1 molecule. (D) Measurements of mDia1/2 elongation velocities *in vivo*. Left to right: Scheme of HT1080 (top) and EL4 (lower) cells, kymographs of single mNeonGreen-mDia1 (left) or mDia2 (right) molecules followed by velocity distributions. Lines are Gaussian fits. Means and SD are indicated. [N_cells_ ≥10, n_molecules/cell_ ≥30, n_total_ ≥650 per condition]. (E) Workflow to generate profilin1 and β-actin overexpressing HT1080 cells. Polycistronic constructs for β-actin, mScarletI and profilin1 were integrated into the genome. Cells were sorted into four sub-populations dependent on mScarletI fluorescence intensity (Fig. S6B, Methods). (F) Top: Western blot of HT1080 cells (wt or overexpressing sub-populations). No translational read-through is visible (1xRP: actin-mScarletI, 2xRP: actin-mScarletI-profilin1 at expected Mw). Bottom: Relative profilin1 and actin levels (fold over wt) for indicated sub-populations. (G) mDia1/2 velocities in profilin-actin overexpressing HT1080 cells. Left to right: Scheme, kymographs of single mNeonGreen-mDia1 (left) or mDia2 (right) molecules, velocity distributions. Lines are Gaussian fits (Red continuous (PA-OE) and dashed (wt) cells as in (D)). Means and SD are indicated. [N_cells_ ≥10, n_molecules/cell_ ≥30, n_total_ ≥650 per condition]. (H) Mean mDia velocities in HT1080 cells plotted against the relative profilin-actin concentration. Error = SD. Dashed lines are linear fits through the origin.

To test for cell-type specific regulation as a reason for this invariance, we perturbed profilin-actin levels in a single cell type. Given their low profilin-actin concentration (Fig. 1D), we overexpressed profilin-actin in HT1080 cells. To prevent side-effects anticipated for the overexpression of profilin alone, we co-overexpressed profilin and actin. To this end, we integrated β-actin with profilin1 and Scarlet-I (as a fluorescent reporter), separated by ribosomal skip sites into a single transgene (Fig. 6E, Methods). We sorted a heterogeneous pool of stably expressing cells into sub-populations depending on reporter fluorescence (Fig. S6B). Quantification showed that balanced profilin-actin overexpression (2-3-fold) could be achieved in the strongest overexpressing subpopulation (Fig. 6F & S6A). We then analyzed the speed of formin-driven actin elongation in these cells. Strikingly, we observed only a marginal increase in mDia1 and mDia2 velocities (by 20 and 12%, respectively) compared to wildtype HT1080 cells (Fig. 6G). Plotting these velocities against the measured relative profilin-actin levels shows that formin-driven elongation neither strongly nor linearly scaled with profilin-actin concentration (Fig. 6H). This was also evident when examining formin velocities from both HT1080 and EL4 cells as a function of the various absolute profilin-actin concentration these cells contained (Fig. S6C). Instead, the data could be well fit with saturation kinetics very similar to those *in vitro* (Fig. S6C). We conclude that formin-mediated actin elongation in mammalian cells i) is resilient to variations in profilin-actin levels and ii) closely matches the maximal *in vitro* rates. These findings combined strongly suggest that mammalian cells maintain profilin-actin concentrations near saturation. More importantly, they also indicate that the kinetic limit to actin filament elongation imposed by profilin and formin we discovered *in vitro* similarly operates in the cytoplasm of living cells.

## Discussion

We have uncovered a biochemical mechanism that controls actin filament growth at physiological subunit levels (Fig. 7). The mechanism provides robustness to actin dynamics, because it buffers filament growth against changes in the concentration of polymerizable actin in different cellular contexts and across cell types. It is based on two central elements: i) A kinetic bottleneck in filament elongation limiting growth and ii) maintenance of profilin-actin concentrations near saturation. Both features emerge from the versatile biochemical activities of profilin.

**Fig. 7:**
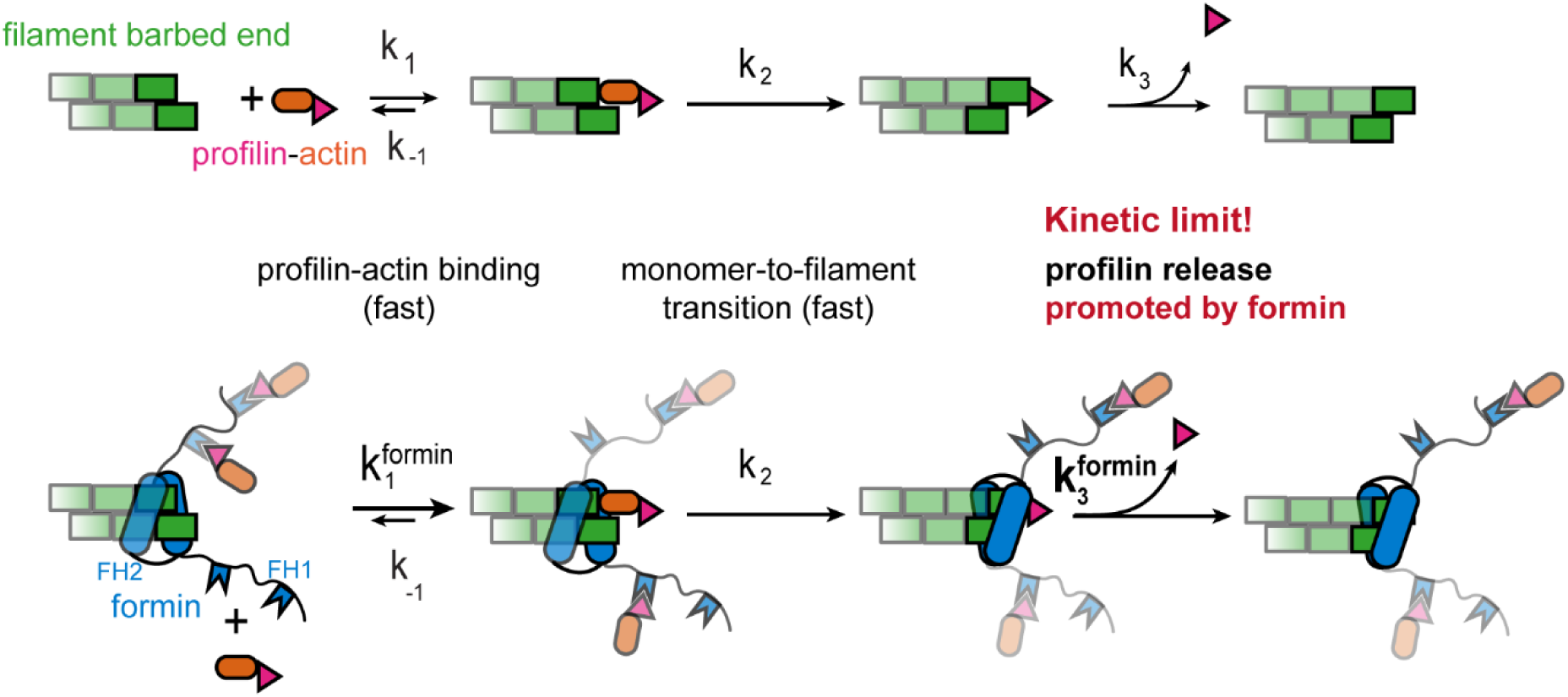
Profilin release controls the speed of actin filament growth. Kinetic scheme of the filament elongation cycle from profilin-actin either in the absence (top) or the presence (bottom) of formins. Reaction 1 and 2 are very fast at physiological profilin-actin concentrations, which is why reaction 3 (profilin release from the terminal protomer) kinetically limits the elongation cycle. Formins accelerate both the first and third reaction of the cycle.

The use of mammalian non-muscle actin in uniquely sensitive assays enabled us to identify the constraints profilin imposes on actin filament growth at physiological conditions. Profilin release is very rapid, but nonetheless kinetically limits filament elongation. The structural monomer-to-filament transition is likely even faster and independent of ATP hydrolysis in actin. This agrees with recent work showing that major structural rearrangements in actin upon polymerization are independent of nucleotide state changes (Merino et al., 2018) and with indirect biochemical evidence (Jégou et al., 2011). Profilin binds very weakly (K_D_ > 20 μM) to ATP-bound filament barbed ends (Courtemanche and Pollard, 2013; Pernier et al., 2016) but very tightly to actin monomers (K_D_ < 0.1 μM). As such, it should out-compete other abundant monomer-binding proteins. The free profilin concentrations required to simply bind barbed ends from solution are thus unlikely attained in mammalian cells. Our results nonetheless imply that many growing filament ends in cells are decorated with profilin, however, not as a result of equilibrium binding but through an active, polymerization-coupled mechanism.

Surprisingly, formins stimulate filament growth even at physiological profilin-actin levels. They do so by promoting profilin release through their FH2 domain (Fig. 7). Profilin and formin appear to mutually destabilize each other at barbed ends, because profilin is known to inhibit formin end-binding (Pernier et al., 2016). Structural models suggest that profilin and the formin FH2 domain might directly interfere at barbed ends (Fig. S5D). Alternatively, formins could alter the end structure (Aydin et al., 2018) to promote profilin release allosterically. Structures of formin- and profilin-bound barbed ends will be required to resolve this question. Whether and how polymerases unrelated to formins such as Ena/VASP proteins also promote profilin release will be important to study in the future.

Do all actin filaments in mammalian cells grow at their maximal, profilin release-limited speed? The growth speeds we observe are faster than actin network movement in many cellular protrusions (Renkawitz et al., 2009), indicating that this is unlikely the case. This mismatch might be explained by filament orientation and, more importantly, compressive forces that slow down filaments pushing against membranes (Bieling et al., 2016; Mueller et al., 2017). Forces might change the rate-limiting step of filament growth, because they will likely not affect all reactions of elongation equally (Fig. 7). Brownian ratchet models predict that compression should strongly inhibit the binding of profilin-actin (Mogilner and Oster, 1996). The profilin-actin concentration might thus still affect the growth of filaments experiencing load and the force they generate. This might explain why rapidly moving cells contain higher profilin-actin levels.

Competition for soluble actin has been proposed (Suarez and Kovar, 2016) to explain mutual inhibition between distinct actin structures in cells (Rotty et al., 2015; Suarez et al., 2015). How can this be reconciled with robust actin growth from a large profilin-actin pool we observe? Mutual inhibition might not necessarily originate from simple competition, but effects that can be understood when taking free actin monomers into consideration. Because the cellular profilin pool is finite and lower than the actin concentration, disassembly of entire network types might transiently exceed profilin’s capacity to bind actin monomers. Since actin nucleation -both catalyzed and spontaneous- is strongly promoted by free actin monomers, this might trigger nucleation through alternative pathways resulting in homeostatic filament amounts. Such a monomer-triggered mechanism has been proposed for formin-mediated nucleation after cell deformation (Higashida et al., 2013). Ways to detect distinct soluble actin states *in vivo* are needed to understand their effect on local actin network dynamics (Skruber et al., 2018).

## Acknowledgements

We thank Philippe Bastiaens for support and help in shaping the manuscript. Scott Hansen, Dyche Mullins, Thomas Surrey, Andrea Musacchio and Stefan Westermann for comments on the manuscript and Marcus Taylor for EL-4 cells. This work was supported by HSFP (CDA00070/2017-2), the MaxSynBio network and the Max Planck Society.

## Author contribution

J. F. and P. B. designed and conducted experiments, analyzed data, and wrote the manuscript, F. M. under supervision of S. R. generated structural models and designed profilin and actin mutants. J. F., L. V., P. V. and M. P. isolated immune cells and performed cell volume measurements.

## Supplementary materials

### Material and Methods

#### Contact for reagent and resource sharing

Further information and requests for resources and reagents should be directed to and will be fulfilled by the Lead Contact, Peter Bieling.

### Methods

#### Structural models of barbed end complexes

Using MODELLER (Webb and Sali, 2016) we built models of the binding of profilin, formin or profilin-formin to the barbed end of the actin filament. For the profilin models we superimposed the actin monomer in the profilin/β-actin crystal structure (PDBID 2BTF) (Schutt et al., 1993) to either the ultimate or penultimate subunit of a filament barbed end. As a filament template we used our recent structure of α-actin in complex with beryllium fluoride (PDBID 5OOF) (Merino et al., 2018).

To model the FH2 domain of mDia1 (aa 750-1163) bound to a profilin-occupied barbed end we superimposed subdomains 1 (aa 1-33; 70-137; 348-375) and 3 (aa 138-180; 274-347) of an actin subunit from the Bni1p-actin crystal structure (PDBID 1Y64) (Otomo et al., 2005) with the terminal monomers in F-actin-BeFx. This brings the FH2 domain of the formin to the right position in the actin filament. Given that the Bni1p structure has a non-physiological helical arrangement of the formin, we erased the loop between its Knob and Lasso regions (aa 804-831 in mDia1) and built it de novo to recover the known dimeric arrangement of the FH2 domains. To further improve the quality of the models we also included two crystal structures of the FH2 domains of mDia1 (PDBID 1V9D and 3O4X) (Nezami et al., 2010; Shimada et al., 2004).

#### Protein design

We used the RossetaScripts framework(Fleishman et al., 2011) within Rosetta(Leaver-Fay et al., 2011) to find possible mutations to increase the affinity of profilin for actin. For the design we tested a model built on the crystal structure of profilin-β-actin (Schutt et al., 1993) as well as our F-actin-profilin models (see previous section). The design strategy was modified from the protocol provided by the Baker lab in (Berger et al., 2016) (see Computational methods: design with ROSETTA in their manuscript). We tried mutating all profilin residues at the interface with actin, but did not allow mutations into Cys, Pro, Trp, or Gly. We generated a total of 1920 possible profilin sequences for each actin conformation, and kept the top 50 (lowest energies) for further analysis. From there, we selected single mutations likely to increase the affinity of profilin for actin and tested them experimentally.

#### Protein purification and labeling

##### 10xhis-Gelsolin G4-6

Mouse Gelsolin G4-6 was cloned with an N-terminal 10xhis tag into a pCOLD vector. Protein was expressed in E. coli BL21 Rosetta cells for 16 hrs at 16 °C. After cell lysis (20 mM Tris-Cl pH 8.0, 300 mM KCl, 5 mM CaCl_2_, 0.2 mM ATP, 0.5 mM β-mercaptoethanol, 1 mM PMSF, DNAseI) the lysate was hard spun and purified by IMAC over a 40 ml Ni^2+^ superflow column. Protein was gradient eluted (20 mM Tris-Cl pH 8.0, 300 mM KCl, 5 mM CaCl_2_, 0.2 mM ATP, 500 mM Imidazole) over 10 column volumes followed by gelfiltration over Superdex 200 26/600 into storage buffer (5 mM Tris-Cl pH 8.0, 50 mM KCl, 5 mM CaCl_2_, 0.1 mM ATP, 0.5 mM TCEP, 20 % glycerol). The protein was snap frozen in liquid nitrogen and placed in −80 °C for long-term storage.

##### Native bovine (β, γ)-actin

Bovine thymus was manually severed into small fragments and mixed in a precooled blender together with ice cold Holo-Extraction buffer (10 mM Tris-Cl pH 8.0, 7.5 mM CaCl2, 1 mM ATP, 5 mM β-mercaptoethanol, 0.03 mg/ml benzamidine, 1 mM PMSF, 0.04 mg/ml trypsin inhibitor, 0.02 mg/ml leupeptin, 0.01 mg/ml pepstatin, 0.01 mg/ml apoprotein). After homogenizing, additional 2.5 mM β-mercaptoethanol was added to the lysate and the pH was checked and readjusted to pH 8.0 if necessary. After initial centrifugation the lysate was filtered through a nylon membrane [100 μm] and hard spun in an ultracentrifuge. The volume of the cleared supernatant was measured out and the salt and the imidazole concentrations were adjusted (KCl to 50 mM, imidazole to 20 mM). The supernatant was incubated with the gelsolin G4-6 fragment to promote the formation of actin:gelsolin G4-6 complexes. To this end, 4 mg of 10xhis-gelsolinG4-6 were added for each g of thymus to the lysate and dialyzed into IMAC wash buffer overnight (10 mM Tris-Cl pH 8.0, 50 mM KCl, 20 mM imidazole, 5 mM CaCl_2_, 0.15 mM ATP, 5 mM β-mercaptoethanol). The lysate containing the actin:gelsolin G4-6 complex was then circulated over a Ni^2+^ superflow column. Actin monomers were eluted with Elution Buffer (10 mM Tris-Cl pH 8.0, 50 mM KCl, 20 mM imidazole, 5 mM EGTA, 0.15 mM ATP, 5 mM β-mercaptoethanol) into a collection tray containing MgCl_2_ (2mM final concentration). Actin containing fractions were identified by gelation, pooled and further polymerized for 4 hrs at RT after adjusting to 1xKMEI and 0.5 mM ATP. After ultracentrifugation, the actin filament pellet was resuspended in F buffer (1xKMEI, 1xBufferA) and stored in continuous dialysis at 4 °C. F buffer containing fresh ATP and TCEP was continuously exchanged every 4 weeks.

For fluorescence measurements actin monomers were labelled with 1.5-IAEDANS at Cys374 as outlined in (Hudson and Weber, 1973; Miki et al., 1987) using a modified protocol. Briefly, the actin filament solution was transferred to RT, mixed with 10x molar excess of 1.5-IAEDANS and incubated for 1 hr at RT. The reaction was quenched by the addition of 1 mM DTT for 10 min. After ultracentrifugation at 500.000xg for 30 min, the actin pellet was resuspended in an appropriate amount of BufferA and dialyzed in the same buffer at 4 °C for 2 days. Actin monomers were separated from residual filaments by centrifugation at 300.000xg followed by determination of monomer concentration and degree of labelling at 280 nm/336 nm.

##### Recombinant human β-actin

Human β-actin was cloned with a C-terminal linker sequence (ASRGGSGGSSGGSA) followed by the human β-thymosin sequence followed by a 10xhis tag (Noguchi et al., 2007) in a pFL vector. PCR based site directed mutagenesis was performed to generate human, ATPase deficient β-actin (Q137A+D154A+H161A). Proteins were expressed in insect TnaO38 cells for 3 days at 27 °C. The cells were resuspended with a 5x pellet volume of lysis buffer (10 mM Tris-Cl pH 8.0, 50 mM KCl, 7.5 mM CaCl_2_, 1 mM ATP, 5 mM imidazole, 5 mM β-mercaptoethanol, 0.03 mg/ml benzamidine, 1 mM PMSF, 1x complete protease inhibitor cocktail). After cell lysis by a microfluidizer the lysate was hard spun, filtered through a 0.45 μm syringe filter and passed through a Ni^2+^-sepharose excel column. After washing the column with 10 mM Tris-Cl pH 8.0, 50 mM KCl, 5 mM CaCl_2_, 0.15 mM ATP, 5 mM imidazole, 5 mM β-mercaptoethanol, the protein was eluted over a 6 CV linear gradient to Elution Buffer (10 mM Tris-Cl pH 8.0, 50 mM KCl, 0.15 mM ATP, 300 mM imidazole, 5 mM β-mercaptoethanol) followed by dialysis into BufferA overnight. Next, the protein was cleaved with TLCK-treated chymotrypsin in a molar ratio of 250:1 (actin over chymotrypsin) at 25 °C. After 10 min the reaction was quenched with 0.2 mM PMSF at 4 °C. The protein was again passed over the Ni^2+^-sepharose excel column and the flow through was polymerized for 3 hrs at 25 °C by the addition of 1xKMEI, 2 mM MgCl_2_ and 0.5 mM ATP. After hard spin, the actin filament pellet was resuspended into F buffer (1xKMEI, 1xBufferA) and stored in dialysis at 4 °C.

##### Profilin 1 and 2

Human profilin isoforms 1 and 2 were expressed either as untagged proteins or with an N-terminal SUMO3-10xhis tag in E. coli BL21 Rosetta cells at 30 °C for 4.5 hrs. Profilin1 mutants that were generated via site directed mutagenesis (E82A, R88K, K125E E129K and S71M) were expressed with an N-terminal SUMO3-10xhis in E. coli BL21 Rosetta cells at 30 °C for 4.5 hrs. For the N-terminal SUMO3-10xhis tagged version, the cells were lysed (20 mM Tris-Cl pH 8.0, 300 mM NaCl, 10 mM imidazole, 0.5 mM β-mercaptoethanol, 15 μg/ml benzamidine, 1 mM PMSF and DNaseI) and the hard spun lysate was circulated over a 5 ml HiTrap Chelating column followed by overnight SenP2 cleavage of the N-terminal SUMO-his tag on the column, generating the natural profilin N-terminus. After cleavage the flow through was gelfiltered over a Superdex 200 16/600 column into storage buffer (20 mM Tris-Cl pH 7.5, 50 mM NaCl, 0.5 mM TCEP). The non-tagged profilin isoforms were purified as described in(Bieling et al., 2018) by ammonium sulfate precipitation, followed by ion-exchange (DEAE) and hydroxylapatite (HA) chromatography steps, followed by size exclusion chromatography (Superdex 200 16/600) into storage buffer (20 mM Tris-Cl pH 7.5, 50 mM NaCl, 0.5 mM TCEP). Proteins were snap frozen in liquid nitrogen with the addition of 20 % glycerol in the storage buffer and were stored at −80 °C

##### Profilin - Actin complex

Filamentous mammalian actin was depolymerized through dialysis into BufferA (2 mM Tris, 0.2 mM ATP, 0.1 mM CaCl_2_, 0.1 μg/ml NaN_3_, 0.5 mM TCEP) and gelfiltered over a Superdex 200 16/600. After gelfiltration a 1.5x molar excess of profilin was added to the actin monomers and incubated at 4 °C overnight to form profilin-actin complexes. Profilin-actin was then separated from excess free profilin by gelfiltration over a Superdex 200 10/300 GL into BufferA. The complex was concentrated to working concentrations between 200-400 μM and stored at 4 °C up to two weeks without inducing nucleation.

##### Formins

*M. musculus* mDia1 FH1-2 (aa 548-1154), mDia2 FH1-2 (aa 515-1013), FH1_mdia1_FH2_mdia2_ (aa 548-751/453-1013), FH1_mdia2_FH2_mdia1_ (aa 515-612/645-1154), *H. sapiens* DAAM1 FH1-2 (aa 490-1029), were expressed with an N-terminal 10xhis-SNAP-tag. All constructs were expressed in E. coli BL21 Star pRARE cells for 16 hrs at 18 °C. The cells were lysed in Lysis Buffer (50 mM NaPO_4_ pH 8.0 (pH 7.5 for mDia chimera constructs), 400 mM NaCl, 0.75 mM β-mercaptoethanol, 15 μg/ml benzamidine, 1xcomplete protease inhibitors, 1 mM PMSF, DNaseI,) and the protein was purified by IMAC using a 5 ml HiTrap column. The protein was eluted using Elution Buffer (50 mM NaPO_4_ pH 7.5, 400 mM NaCl, 400 mM imidazole, 0.5 mM β-mercaptoethanol) in a gradient and the 10xhis-tag was directly cleaved using TEV protease overnight. After cleavage proteins were desalted into low salt Mono S buffer (10 mM Hepes pH 7.0 (pH 7.5 for mDia chimera constructs), 90 mM NaCl, 0.5 mM TCEP) over a HiPrep 26/10 desalting column followed loading onto a MonoS column. Protein was eluted by a linear 25 column volume gradient to high salt MonoS buffer (10 mM Hepes pH 7.5, 1 M NaCl, 0.5 mM TCEP) followed by gelfiltration over a Superdex 200 16/600 into storage buffer (20 mM Hepes pH 7.5, 200 mM NaCl, 0.5 mM TCEP, 20 % glycerol).

Following the purification the proteins were either snap frozen and stored in –80 °C or directly used for SNAP-labeling. A 3x molar excess of SNAP Cell TMR-star was mixed with the protein and incubated for 6 hrs at 16 °C followed by an overnight incubation on ice. Post labeling the protein was gelfiltered over a Superose 6 10/300 GL column into storage buffer. The degree of labelling (50–70 %) was determined by absorbance at 280 nm and 554 nm.

##### Myosin and biotinylated heavy - mero – myosin (HMM)

Skeletal muscle myosin was prepared from chicken according to(Pollard, 1982). Briefly, 300 g muscle tissue were mixed with 4x volumes of extraction buffer (0.15 mM KH_2_PO_4_ pH 6.5, 0.3 M KCl, 5 mM MgCl_2_, 0.1 mM ATP, 20 mM EDTA) while blending. The pH was adjusted to 6.6 afterwards. After centrifugation, the supernatant was diluted with 10x volumes of cold water and the precipitate was separated from solution by centrifugation at 9.000xg for 30 min. The pellet was resuspended in buffer 8 (3 ml buffer per g of pellet, 60 mM KH_2_PO_4_ pH 6.5, 1 M KCl, 25 mM EDTA) and dialyzed against buffer 9 (25 mM KH_2_PO_4_ pH 6.5, 0.6 M KCl, 10 mM EDTA, 1 mM DTT) over night. Following dialysis, an equal volume of cold water was added to the myosin solution and stirred for 30 min. After centrifugation for 30 min at 15.000xg, the supernatant was diluted with 7 volumes of cold water and again spun for 30 min at 9.000xg. The pellet fraction was then resuspended into buffer 10 (20 mM Tris-Cl pH 7.0, 0.6 M KCl, 10 mM DTT) and treated with α-chymotrypsin (25 μg/ml final) at 25 °C for 15 min. The reaction was quenched by the addition of 0.3 mM PMSF. After protease treatment, the myosin was dialyzed into buffer 11 (10 mM NaPi pH 7.2, 35 mM NaCl, 10 mM DTT). On the next day, the HMM was separated by ultra-centrifugation for 1 hr at 300.000xg. The supernatant was desalted into buffer 11 without DTT and incubated with 15x molar excess of EZ-Link maleimide-PEG11-biotin for 2 hrs on ice. The reaction was stopped by the addition of 1 mM DTT. The protein was desalted into buffer 11 containing 20 % glycerol, SNAP-frozen and stored at –80 °C.

#### Endpoint hydrolysis measurements via HPLC

All HPLC measurements were initiated by loading actin monomers and profilin-actin with Mg-ATP. After a 1 hr incubation of monomers and profilin-actin (40 μM) with 1 mM MgCl_2_ and 1 mM ATP, proteins were desalted into 2 mM Tris-Cl pH 8.0 using a Zeba Spin Desalting column. Actin seeds were then polymerized from the desalted actin monomers by adjusting to 1xKMI (50 mM KCl pH 7.0, 1.5 mM MgCl_2_, 10 mM imidazole) for 1 hr at 23 °C. To start the reaction, profilin–actin (40 μM) was mixed with seeds (5 μM) in presence of 1xKMI. After 1.5 hrs incubation at 23 °C, the samples were boiled for 5 min followed by a hard spin. The supernatant was carefully aspired and analyzed by HPLC. As a negative control, profilin-actin were stabilized with 5 mM latrunculin B and the seeds were incubated with 5 mM phalloidin before mixing, otherwise the samples were treated as mentioned above.

All nucleotide retention times were measured using an UltiMate 3000 HPLC Dionex – System. The samples were injected onto a C18-column equilibrated with 16 % acetonitrile, 50 mM KP_i_ pH 6.6, 10 mM TBABr. The nucleotide signal intensity was recorded at 254 nm.

#### Radioactive ATPase assays

100 μM Mg-ATP-actin was dialyzed into BufferA for 7 days. After gelfiltration over a Superdex200 16/60 the actin monomer fraction was split into two fractions. With the addition of 1.5x-molar excess profilin1 to one of the monomer fractions, profilin-actin complexes were formed and isolated over a Superdex200 10/300 GL. Both actin monomer and profilin-actin fractions were desalted into ATP free BufferA (2 mM Tris-Cl pH 8.0) over a Zeba Desalting column. 1 ml of 10 μM actin monomers was incubated with 2xKMEI to polymerize actin for 1 hr at RT. In the meantime, 1 ml of 10 μM profilin-actin was incubated with 0.2 mM EGTA, 0.132 mM MgCl_2_ and 0.06 mM γ–^32^P–ATP (3000 Ci/mmol, PerkinElmer #NEG002A) for 30 min on ice. After incubation, γ– ^32^P–ATP labeled profilin-actin complexes were desalted over a Zeba Desalting column into 2 mM Tris–Cl pH 8.0, 0.2 mM EGTA, 0.132 mM MgCl_2_. Immediately before introducing the pre-polymerized actin seeds into the experiments, seeds were sheared through a 27 G needle. The ATPase assay reaction was started by rapidly mixing 6 μM of actin seeds with 6 μM of γ–^32^P–ATP labeled profilin-actin. 100 μl samples were taken at different time points over a time course of 48 min and immediately quenched with an equal volume of silicotungstic–sulfuric acid (4.3 % aqueous silicotungstic acid in 2.8 N sulfuric acid). Samples were recovered in 1 ml of a 1:1 isobutanol/xylene solution and immediately rigorously mixed with additional 100 μl of 10 % ammonium molybdate for 20 s. After 4 min centrifugation at 200xg the upper phase containing the phosphate molybdate complex was extracted. The complex was diluted in LSC cocktail (Hidex) and the number of counts was detected using a liquid scintillation counter (Triathler multilabel tester, Hidex).

#### Fluorescence anisotropy experiments

The measurements were performed in 96 well CORNING plates with a TECAN SPARK plate reader. A constant concentration of 150 nM actin monomers were stabilized with 25 μM latrunculin B and mixed with 4 nM Atto488-WAVE1(WCA) (Bieling et al., 2018). Profilin was titrated to the Atto488-WAVE1(WCA):actin complex to final concentrations of 0–20 μM and equilibrated for 5 min at RT before the measurement. The assay was performed in 1xTIRF buffer (20 mM Hepes pH 7.0, 100 mM KCl, 1.5 mM MgCl_2_, 1 mM EDTA, 20 mM β-mercaptoethanol, 0.1 mg/ml β-casein, 1 mM ATP). For the determination of anisotropy values, Atto488-WAVE1(WCA) was excited at 485/20 nm and the emission was detected at 535/25 nm.

#### IAEDANS fluorescence quenching measurements

Fluorescence measurements were performed in 96 well CORNING plates with a TECAN SPARK plate reader. A constant concentration of 150 nM 1.5-IAEDANS labelled actin monomers were pre-mixed with 25 μM latrunculin B in 1xTIRF assay buffer and thymosin-β_4_ was titrated over a range of 0–200 μM. The 1.5-IAEDANS actin was excited at 336 nm and the emission and thus the fluorescence change of the 1.5-IAEDANS actin bound to thymosin-β_4_ was detected at 490 nm.

#### Tryptophan fluorescence quenching by stopped flow

To determine the association rate constant for profilin binding to actin monomers, increasing profilin concentrations were mixed in a 1:1 volume with a fixed concentration of 0.5 μM actin monomers at 25 °C. The assay was performed in 20 mM Hepes pH 7.0, 100 mM KCl, 1.5 mM MgCl_2_, 1 mM EDTA, 20 mM β-mercaptoethanol, 1 mM ATP, 1.5 μM latrunculin B. Tryptophan fluorescence intensity was recorded by a SX20 double mixing stopped flow device (Photophysics) using excitation and emission wavelengths of 280 and 320 nm, respectively. The time courses of tryptophan fluorescence was recorded and fitted with a single exponential function to yield the observed pseudo-first order reaction rate (k_obs_) as a function of profilin concentration.

#### Single filament experiments on functionalized glass coverslips using TIRF-Microscopy

Flow chambers were prepared from microscopy counter slides passivated with PLL-PEG and coverslips (22×22 mm, 1.5 h, Marienfeld-Superior) that were functionalized according to(Bieling et al., 2016). Briefly, coverslips were cleaned with 3 M NaOH and Piranha solution followed by silanization and PEG-biotin/hydroxy functionalization. For the single filament assays the flow cell surfaces were blocked for 5 min with a Pluronic block solution (0.1 mg/ml κ-Casein, 1 % Pluronic F-127, 1 mM TCEP, 1xKMEI), followed by 2 washes with 40 μl of wash buffer (0.5 mM ATP, 1 mM TCEP, 1xKMEI, 0.1 mg/ml β-Casein). The channel was incubated with 75 nM streptavidin for 3 min, followed by washing and incubation of 90 nM biotin-phalloidin for 3 min. Pre-polymerized actin seeds were immobilized in the channel for another 2 min for cases when spontaneous nucleation was not rapid enough (e.g. low profilin-actin concentrations, absence of formins).

Visualization by TIRF-M was performed following a modified protocol as outlined in (Hansen and Mullins, 2010; Kuhn and Pollard, 2005). Briefly, 9 μl of a 4.44x μM profilin-actin solution was mixed with 1 μl of 10x ME (0.5 mM MgCl_2_, 2 mM EGTA) and 4 μl oxygen scavenging system (1.25 mg/ml glucose-oxidase, 0.2 mg/ml catalase, 400 mM glucose) (Aitken et al., 2008; Bieling et al., 2010; Rasnik et al., 2006). The Mg-ATP–profilin-actin was then combined with 26 μl reaction buffer mix containing additives including 10 nM Cy5-UTRN_261_, (plus additives as described in the specific results section and in the corresponding figure legends) and TIRF buffer with the final composition of: 20 mM Hepes pH 7.0, 100 mM KCl, 1.5 mM MgCl_2_, 1 mM EDTA, 20 mM β-mercaptoethanol, 0.1 mg/ml β-casein, 0.2 % methylcellulose (cP400, M0262, Sigma-Aldrich), 1 mM ATP and 2 mM Trolox.

Filaments that appeared to either stop growing due to surface defects or that showed very large movements out of the TIRF field were not analyzed. All single filament polymerization experiments were performed using profilin-actin as a substrate unless otherwise indicated in the figure legends.

#### Microfluidic single filament experiments by TIRF microscopy

Experiments were essentially conducted as described in the previous section with the following modifications: Microfludic PDMS chambers were mounted on PEG – biotinylated glass cover slips via plasma treatment as described in (Duellberg et al., 2016). The chambers were designed with 2 or 3 inlets and 1 observation channel. After pluronic block (0.1 mg/ml κ-Casein, 1 % Pluronic F-127, 1 mM TCEP, 1xKMEI) for 5 min, biotinylated Alexa647-phalloidin stabilized actin seeds were bound to the surface via streptavidin. To start actin filament polymerization, profilin-actin was diluted in TIRF buffer and directly transferred from a syringe pump into the reaction chamber to visualize filament elongation immediately under the TIRF-microscope. The flow speed was set to 14-16 μl/min.

#### TIRF-Microscopy data acquisition

All in vitro experiments were performed at RT using a custom built TIRF microscope (OLYMPUS IX81). Image acquisition was done by a EM CCD Andor iXon 888 camera controlled by Micromanager 1.4 software (Edelstein et al., 2014). Fiji ImageJ was used for image and data analysis. Dual color imaging was performed through a 60x OLYMPUS APO N TIRF objective using TOPTICA IBeam smart 640s and 488s/or OBIS 561nm LS lasers and a Quad-Notch filter (400-410/488/561/631-640). Shutters, optical filters, dichroic mirrors and the Andor camera were controlled by Micromanager 1.4 software(Edelstein et al., 2014). Images were acquired between intervals of 0.14 – 10 s using exposure times of 30 – 200 ms to avoid bleaching.

All in vivo single molecule experiments were performed at 23 °C unless otherwise specified using a customized Nikon TIRF Ti2 microscope and Nikon perfect focus system. Image acquisition was achieved by dual camera EM CCD Andor iXon system (Cairn) controlled by NIS – Elements software. Dual color imaging was performed through an Apo TIRF 60x oil DIC N2 objective using a custom multilaser launch system (AcalBFi LC) at 488 nm and 560 nm. Images were acquired at intervals of 0.075 – 0.15 s.

#### Cell culture

HT1080 cells were cultured in DMEM and supplemented with 2 mM glutamine, 1 % NEAA and 10 % FBS. B16F10 cells were cultured in DMEM and supplemented with 4 mM glutamine, 1 % NEAA and 10 % FBS. Mouse EL4 cells were cultured in RPMI-1640 with 10 % FBS. The cells were cultivated at 37 °C with 5 % CO_2_ in a humidified incubator. BMDCs were cultured according to (Vargas et al., 2016). Mouse neutrophil cells were extracted from mouse blood.

#### Quantitative western blot analysis

Quantitative western blots were performed using 12 % SDS gels. To determine actin and profilin amounts per cell, purified actin and profilin references of known concentration were titrated into 1xPBS on the same gel as the cell lysate samples. The number of cells was counted by a Vi-CELL Viability Analyzer from Beckmann Coulter. Cells were lysed in 5 mM Tris–Cl pH 7.5, 150 mM NaCl, 1 mM EDTA, 1 % Triton X-100 and 10 min of sonication. All protein samples were prepared in 1x Laemmli sample loading buffer (Cold Spring Harbor Protocols, 2007). Precision Plus Protein Standard All Blue (Biorad) was used as a molecular weight marker. SDS Gel electrophoresis was performed in Tris-Glycine buffer and proteins were transferred onto a PVDF membrane (Merck Chemicals). After protein transfer membranes were blocked with Odyssey TBS blocking solution (LI-COR Biosciences) for 1 hr at RT and probed with one of the following antibodies: monoclonal mouse anti – actin (1:1000, **#**MA5-11869 ThermoFisher) / profilin1 (1:20000, #061M4892 Sigma) / profilin2 (1:20000, **#**sc-100955 Santa Cruz) and monoclonal rabbit anti - GAPDH(14C10) (1:5000, #2118) as primary antibodies. As secondary antibody infrared labeled - donkey anti-mouse and donkey anti-rabbit were used (1:10000, #925-32212, #926-68073 LI-iCOR). All antibodies were incubated for 1 hr at RT and the membrane was washed with TBS-T (TBS + 0.05 % Tween20) in between. The antibody signal was visualized by fluorescence detection on a LI-COR Odyssey CLx imaging system.

#### Cell volume measurements fluorescence eXclusion

Cell volumes were determined for different cell lines and primary cells as outlined in the text. Measurements were performed as described in (Cadart et al., 2017) for all cell types in suspension or attached to a glass surface using fibronectin.

#### Single molecule visualization of formins in cells

Constitutively active fragments of mDia1 FH1-2 (aa 548-1154) and mDia2 FH1-2 (aa 515-1013) were cloned with an N-terminal mNeonGreen sequence in a pΔCMV vector.

20.000 cells of HT1080 were seeded into a well of an 8 well Lab-Tek 1.5H that was coated with fibronectin (40 μg/ml). On the next day, 1.5 μl FuGENE (Promega) were incubated in 150 μl OptiMEM (Gibco) for 5 min at 23 °C followed by a 15 min incubation with 0.5 μg DNA. The entire transfection mix was directly transferred to the cells.

2×10^6^ EL4 cells were resuspended in 100 μl Nucleofector solution and 2 μg DNA and electroporated by Lonza Amaxa NUcleofector II with the appropriate program. After electroporation, the cells were transferred into 1.5 ml medium. To minimize the transfer of cell debris, cells were once passaged on the following day. Finally, the cells were seeded onto a mouse ICAM-1 coated Lab-Tek 1.5H.

For either cell type after 18 hrs after transfection (HT1080) or initial passage (EL4), the cell culture medium was replaced by HBSS (PAN Biotech #P04-32505). To obtain a more direct comparison with our *in vitro* measurements, which were carried out at room temperature, we imaged cells at room temperature quickly after transferring them to the microscope. Only cells with very low formin expression (<25 molecules per cell per image) were chosen for image acquisition. To prevent an influence of mechanical resistance on formin movement, we only analyzed molecules that translocated freely in the interior of the cell and did not get close the cell periphery, where their movement might be obstructed by the plasma membrane.

Control experiments were performed incubating the cells with either 500 nM latrunculin B, 10 μM Y-27632 or 8 μM JASP (Peng et al., 2011). Imaging was performed either immediately before or 10 min after drug treatment.

#### Overexpression of profilin1 and β - actin in HT1080 cells

Polyclonal HT1080 cell lines were generated using the PiggyBac system according to System Bioscience protocols. For profilin-actin overexpression, the following sequences were cloned in a pBP-CAG vector: human β-actin–P2A–mScaletI–T2A–human profilin1 via Gibson assembly.

After transfection of a construct containing the sequence: actin-P2A-mScarletI-T2A-profilin1, transgenetic cells were selected using puromycin (1 μg/ml) followed by cell sorting through a flow cytometer (BD FACSAria). The distinct sub-populations of the cells were sorted according to their fluorescence intensity and then grown separately. Quantitative western blot analysis were performed to measure the profilin1 and β-actin amounts in these distinct cell populations. We did not detect any actin-containing proteins of larger molecular weight that could potentially result from ribosomal read-through (Fig. 6F), presumably because of actin’s stringent folding requirements.

#### Quantification and statistical data analysis

All analyzed data was plotted and fitted in Origin9.0G. All microscopy experiments were analyzed in ImageJ either manually via kymograph analysis or automated by using the TrackMate plugin (Tinevez et al., 2017) unless otherwise described.

##### Profilin binding affinity for actin monomers by fluorescence anisotropy competition experiments

To determine the equilibrium dissociation constant of profilin (wt or mutant proteins) and actin monomers from competition with another protein (the WCA domain of WAVE1) that binds to actin monomer with known affinity, the mean anisotropy values were plotted against the increasing total profilin concentration [nM]. Mean values were calculated from at least three measurements in three individual experiments per condition, error bars demonstrate the SD. The anisotropy data was fitted by an competitive binding model as described in (Wang, 1995) that analytically solves for the concentrations of the bound and free species from the known total concentrations of all proteins and the equilibrium dissociation constants for each of the two competing ligands:

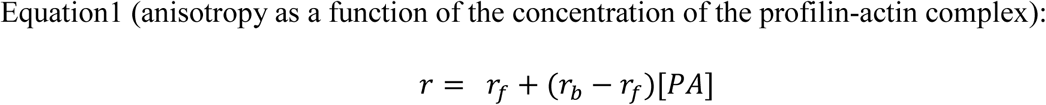

The concentration of the profilin-actin complex can be determined from:

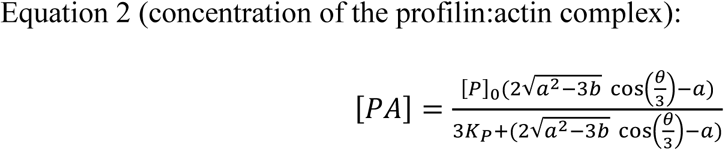

and

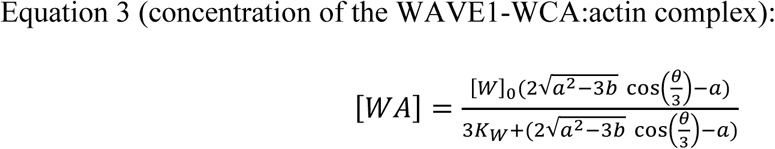

with

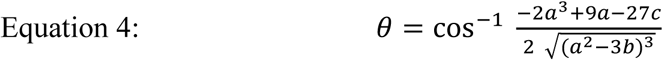

and

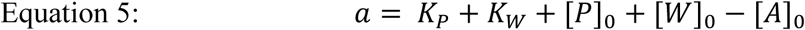

and

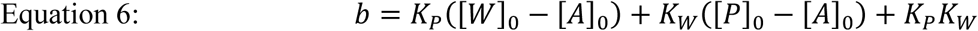

and

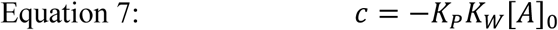

with [A]_0_ being the total actin concentration, [P]_0_ the total concentration of profilin, [W]_0_ the total concentration of Atto488-WAVE1(WCA,) K_P_ the equilibrium dissociation constant for the interaction between profilin and actin and K_W_ the equilibrium dissociation constant for the interaction between Atto488-WAVE1(WCA,) and actin.

##### Thymosin-β_4_ binding affinity for actin monomers by fluorescence measurements

To determine the equilibrium dissociation constant of thymosin-β_4_, the mean decrease in fluorescence intensity [au] was plotted against the increasing total thymosin-β_4_ concentration. Mean values were calculated from at least three measurements in three individual experiments per condition, error bars demonstrate the SD. This data was fitted to a quadratic binding model as described in(Zalevsky et al., 2001):

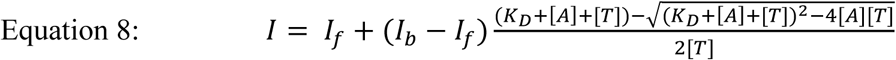

With [A] being the total concentration of IEDANS-labeled actin, [T] the total concentration of thymosin-β_4,_ I_f_ and I_b_ the fluorescent intensities in the free and bound state, respectively and K_D_ being the equilibrium dissociation constant.

##### Calculations of free species

To calculate the free actin, profilin, thymosin-β_4_ (if added) and profilin-actin complex concentrations from the total concentration of actin, profilin and (if added) thymosin-β_4_ in our TIRF-M single filament assays (see Fig. S2B), we used an exact two species competition model as described in(Wang, 1995) and above (see Profilin binding affinity for actin monomers by fluorescence anisotropy competition experiments).

##### Stopped flow measurements

For the determination of the association rate constant for profilin binding to actin monomers by tryptophan fluorescence quenching, the decrease in tryptophan fluorescence [au] was plotted against the total profilin concentration [μM]. The data was fitted with the following mono-exponential decay function to determine k_obs_:

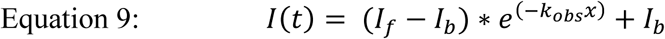

With I(t) the measured fluorescent intensity over time, I_f_ and I_b_ the tryptophan fluorescence in the free and bound state respectively and k_obs_ being the observed reaction rate.

The association rate constants (k_on_) were determined from linear regression fits of the k_obs_ values as function of the total profilin concentration. The dissociation rate constants (k_off_) were calculated from equilibrium dissociation constants (K_D_) and association rate constants (k_on_) using the following equation:

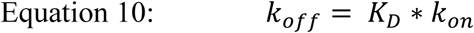

Errors for the dissociation rate constants were calculated using error propagation.

##### Quantitative western blot analysis for profilin and actin

Actin and profilin protein amounts per cell were quantified by western blot analysis using fluorescently-labelled secondary antibodies using a Odyssey Imaging System (LI-COR Biotechnology). The fluorescence signal intensity of the protein bands was analyzed from membrane scans using ImageJ. First, the detected intensity area was selected with the *rectangular tool*, for each protein intensity band (cellular protein and reference protein) an equal sized area was selected. Next, all lanes were plotted in an intensity plot profile reflecting the pixels across the selected area using the command *plot lanes*. The background signal intensity was subtracted from the protein intensity profile by drawing a straight baseline through the intensity curve representing the background intensity to the left and right of the curve. Then, the signal intensity (represented as the area under the intensity profile) was measured by selecting the *tracing tool* and clicking anywhere under the curve to integrate the intensity signal of the area of the plot profile. The measured intensities of the reference protein samples were plotted against the loaded protein mass [ng] and fitted with a linear function. The mass of the protein of interest was then calculated based on the slope of the reference protein. Finally, the protein concentration of actin/profilin was calculated as follows:

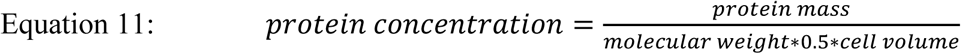

We assumed only half of the total cellular volume because actin and profilin are excluded from the endomembrane system (ER, Golgi, Mitochondria etc.) that occupies roughly 50 % of the cell as measured by tomography methods. This means that in the most extreme case (all of the cell volume can be explored by profilin/actin), we are overestimating protein concentration by maximally 2-fold.

##### Cell volume measurements by fluorescence eXclusion

Data analysis was performed using custom written codes for MATLAB 2017b software written by QuantaCell. First, the raw GFP images were normalized following a manual cell tracking as it has been described earlier from (Cadart et al., 2017). For each cell type we analyzed ≥ 300 single cells. The cell volume distribution was plotted as a histogram and a lognormal distribution curve was fitted to the histogram. The mean volume [μm^3^] and the error (SD) for each cell type was calculated.

##### Barbed end elongation velocity from single filaments by TIRF-microscopy

Images were analyzed by manual filament tracking using the *segmented line tool* from ImageJ and further analyzed by the *kymograph plugin*. The slopes were measured to determine the polymerization rate of individual actin filaments. The pixel size/length was converted into microns/s. One actin monomer contributes to 2.7 nm of the actin filament length. For each experimental condition, the filament polymerization velocity was measured from ≥40 filaments from 3 independent experiments per condition and are reported as mean values with error bars representing SD. The elongation speed as a function of the total profilin-actin concentration were fitted by a hyperbolic model:

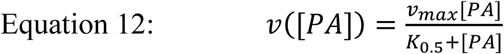

With [PA] being the total profilin-actin concentration, v_max_ the maximal filament polymerization velocity at saturated profilin-actin concentrations and K_0.5_ the profilin-actin concentration at half-maximal elongation speed.

##### Velocity of single formin molecules in vivo

Data analysis was performed by manual filament tracking with the *segmented line tool* from ImageJ. Further, slopes from kymographs were measured to determine the moving rate of individual formins. The pixel size/length was converted into microns/s. One actin monomer contributes to 2.7 nm of the actin filament length. For each experimental condition ≥10 cells and ≥35 single molecules per cell were analyzed. Total number of molecules analyzed per condition was ≥650. All mean speed values were plotted as a histogram and fitted with a Gaussian function.

## Supplementary Figures

**Fig. S1:**
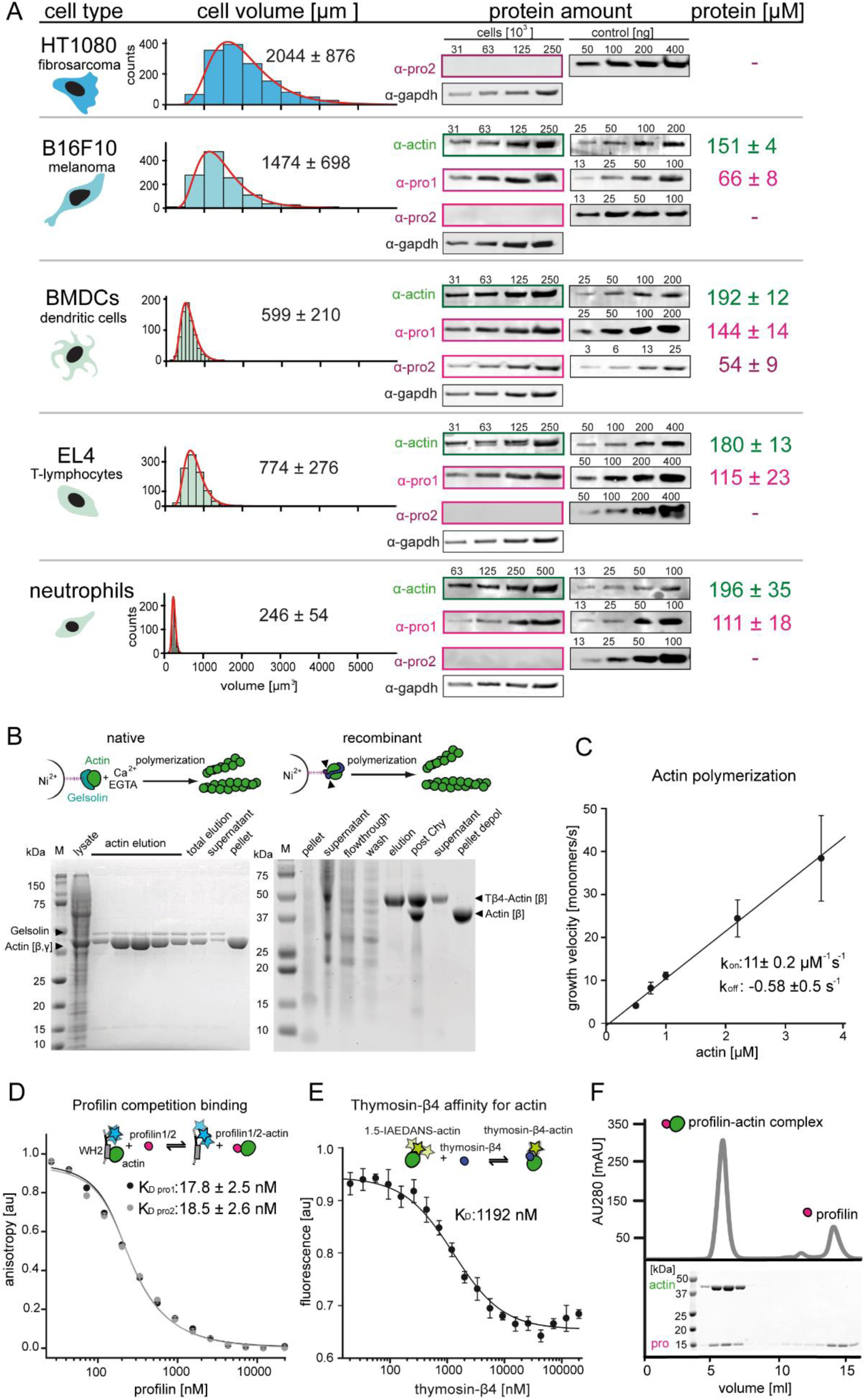
Quantification of profilin-actin levels and purification of mammalian profilin-actin. (A) Profilin-actin concentration determination in mammalian cells. Left to right: name of cell type and origin, histograms of the single cell volume from fluorescence eXclusion measurements, quantitative western blot analysis of cellular actin, profilin1 and profilin-2 amount (left: titration of cell number, right: standard curve of recombinant proteins), calculated mean protein concentration [μM] per cell with experimental error (SD, N=3 independent experiments, see Methods). (B) Purification of mammalian cytoplasmic actin from two sources. Top: Schematic workflow of cytoplasmic actin purification. Left: native purification of β,γ–actin by gelsolin affinity chromatography. Right: Isolation of β–actin from recombinant expression of β–actin–linker-thymosin β_4_ -10xhis and purification via IMAC followed by chymotrypsin cleavage (▴). For both strategies, finally released monomers were polymerized and separated from contaminants. Both purification protocols result in very pure and high yields of protein (see last pellet fraction). (C) Barbed end polymerization rate of cytoplasmic native mammalian (β,γ)–actin as a function of the actin monomer concentration. The mean values ± SD were fitted with a linear function. (D) Binding of profilin1 and 2 to cytoplasmic actin measured by fluorescence anisotropy competition assays. Fluorescence anisotropy of Atto488-WAVE1_WCA_ [4 nM] as a function of increasing profilin1 and 2 concentration in the presence of a constant amount of actin monomers [150 nM]. Lines are fits to an exact analytical competition model (Wang, 1995). Each data point represents the mean value from three independent experiments. Error indicators are SD. (E) Binding of thymosin-β_4_ to cytoplasmic actin measured by fluorescence change assays. Fluorescence measurement of 1.5-IAEDANS labeled actin monomers [150 nM] as a function of increasing thymosin-β_4_ concentration in the presence of a constant amount of actin monomers [150 nM]. Lines are fits to an quadratic binding model (see Methods). Each data point represents the mean value from three independent experiments. Error indicators are SD. (F) Isolation of stoichiometric profilin-actin complexes from free profilin by size exclusion chromatography.

**Fig. S2:**
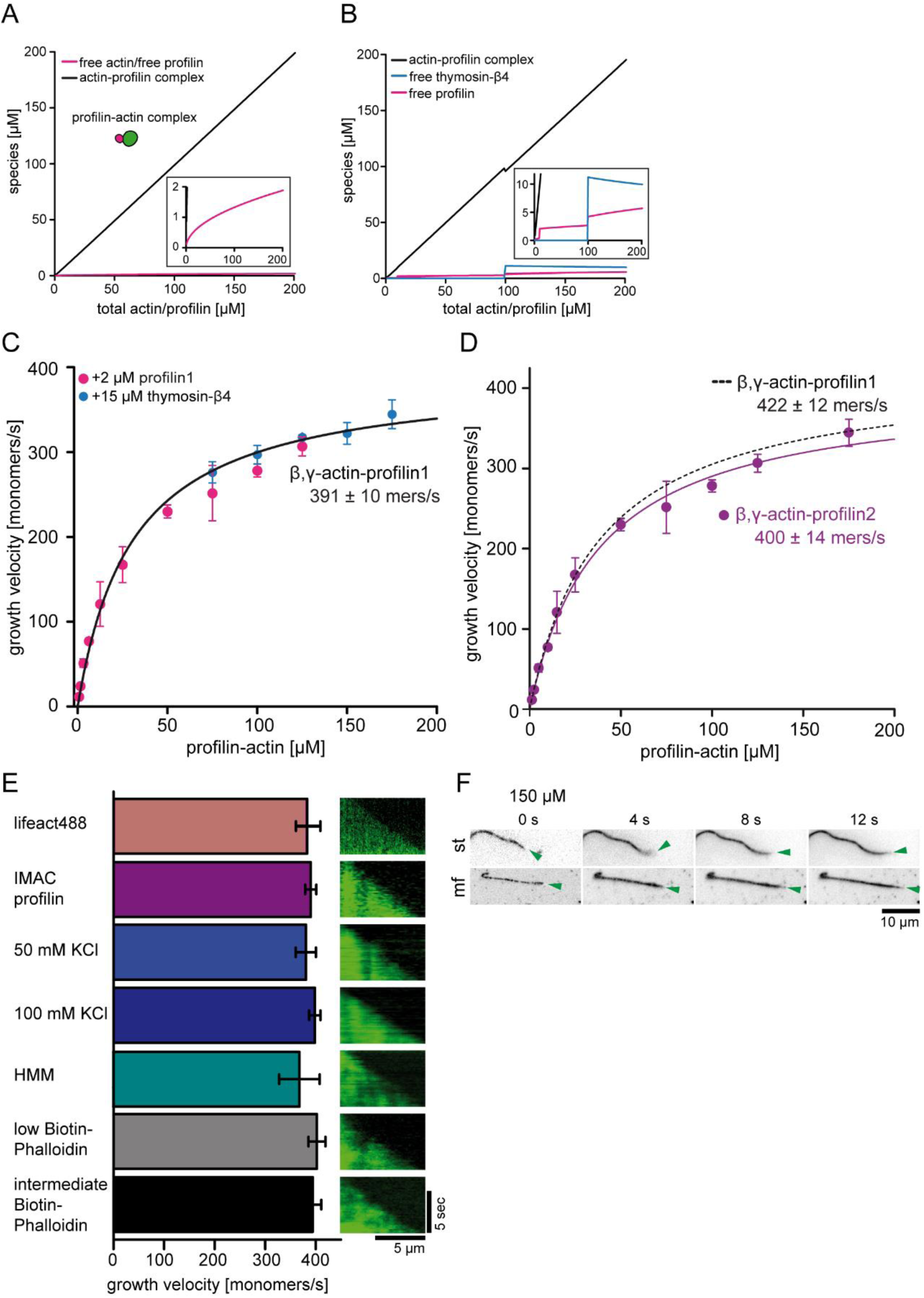
Control experiments for barbed end polymerization in TIRF-M single filament assays. (A) Calculations of profilin-actin complex and free profilin and actin concentrations [μM] (see inset) as a function of the total profilin-actin concentration (see Methods). (B) Calculations of profilin-actin complex and free profilin or thymosin-β_4_ concentrations [μM] by the addition of additional trace amounts of 2 μM profilin (between 10-100 μM total profilin-actin) or 15 μM thymosin-β_4_ (>100 μM total profilin-actin) (see Methods) to suppress residual spontaneous nucleation. Note that no more than 5 μM free profilin or 11 μM free thymosin-β_4_ accumulate in the assay. These low amounts do not significantly affect barbed end growth rates. (C) Barbed end growth velocities measured from TIRF-M single filament assays using profilin1: β, γ–actin as a substrate. In addition 2 μM free profilin1 (magenta) or 15 μM thymosin-β_4_ (blue) at individual and overlapping profilin-actin concentrations indicated in the graph were added to the reaction. Each point represents the calculated mean of the actin filament elongation rate at a distinct substrate concentration [N ≥30 for each condition, error bars = SD]. Continuous line represents a hyperbolic fit yielding the indicated maximum filament growth rate at saturation. (D) Barbed end growth velocities measured from TIRF-M single filament assays using profilin2: β, γ–actin as a substrate (violet). Data from profilin1: β, γ–actin are shown as a dashed black line. Each point represents the calculated mean of the actin filament elongation rate at a distinct substrate concentration [N ≥40 for each condition, error bars = SD]. Continuous lines are hyperbolic fits yielding the indicated maximum filament growth rates at saturation. (E) Controls experiments showing that the maximal filament growth velocity does not depend on the filament binding probe, the profilin purification method, ionic strength or surface attachment. Left: Bar diagram of the growth velocities for fluorescent filament probes (upper 1^st^ - lifeact488, others - Cy5UTRN_261_), affinity purified profilin (upper 2^nd^ – IMAC purified, others – native purification), different salt concentrations (50 and 100 mM KCl), different surface tethering (HMM, biotin-phalloidin low- 20 nM and intermediate- 200 nM) at 100 μM profilin-actin. Right: Kymographs of the growth of representative individual filaments at 100 μM profilin-actin. (F) TIRF-M time-lapse images of filaments either surface tethered along their length (st) or only via a stabilized seed at their pointed end in microfluidic devices (mf). The green arrow follows a single growing barbed end. Experiments were done at 150 μM profilin-actin.

**Fig. S3:**
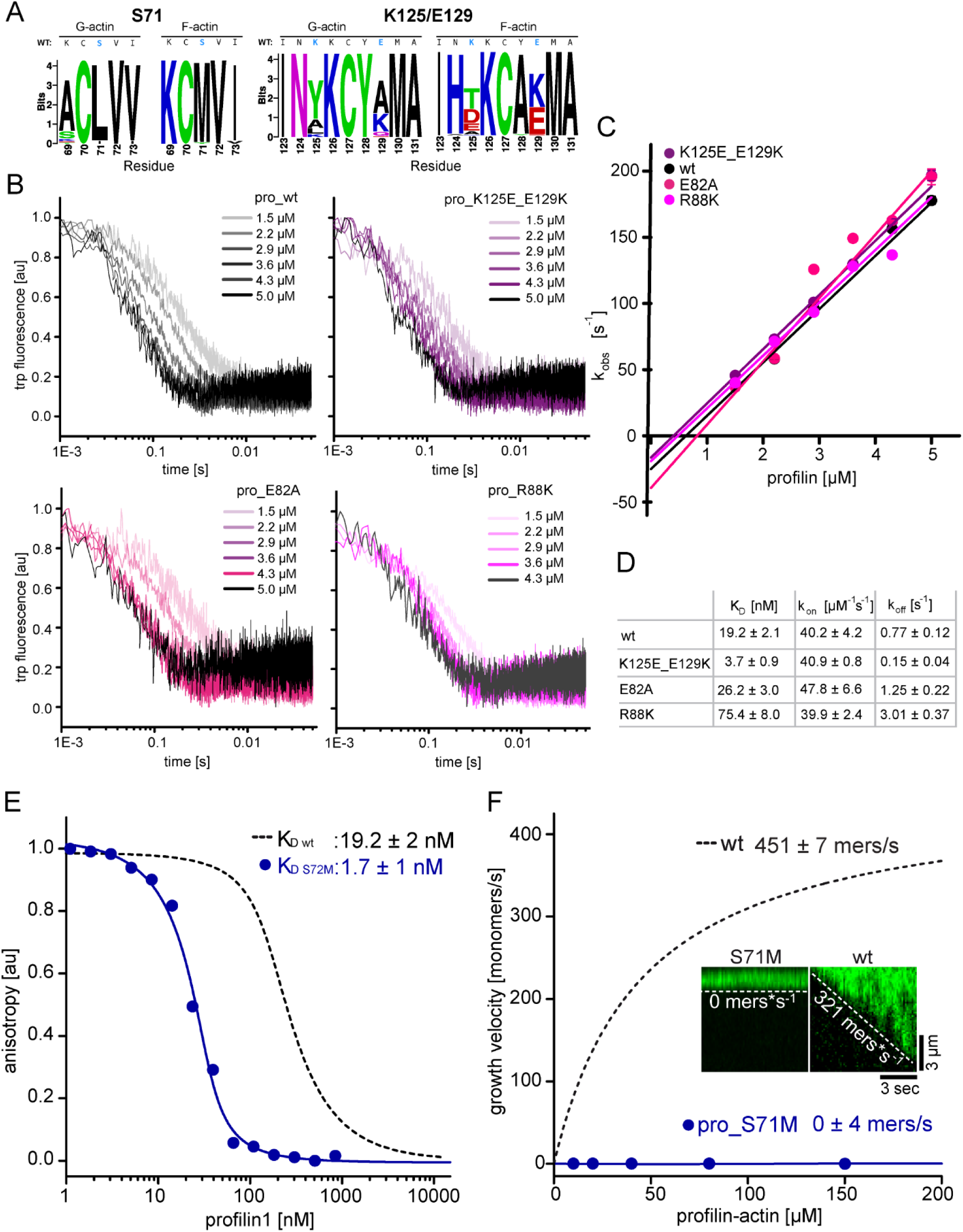
Measurements of profilin1-actin monomer association kinetics and characterization of a ultra-tight binding profilin1 mutant that blocks filament polymerization. (A) Logo for the top 50 designed sequences of profilin. The plot shows the amino acids around the K125/E129 or S71 obtained from the G- and F-actin complexes. For both cases, the wild-type profilin sequence is shown at the top, with the mutated amino acids highlighted in blue. The logos were built with WebLogo (Crooks et al., 2004). (B) Time traces of tryptophan fluorescence quenching upon formation of the profilin1-actin complex for either wt or mutant profilin-1 as indicated. Experiments were done at 0.5 µM actin monomers and various excess profilin-1 concentrations as indicated. The observed reaction rates (k_obs_) (plotted in S3C) were derived from fits to a mono-exponential decay function (see Methods). (C) Linear fit of the observed reaction rates k_obs_ as a function of the total profilin-1 concentration (see Fig. S3B). The association rate constant (k_on_) is calculated from the slope of a linear fit to the data. (D) Summary table of equilibrium dissociation constants (K_D_), association rate constants (k_on_) and dissociation rate constants (k_off_) of the interaction of profilin1 (wt or mutants as indicated) and actin monomers. Dissociation rate constants were calculated from the equilibrium dissociation constants and the measured association rate constants (see Fig. S3B-C, see Methods). (E) Binding of profilin1 (wt and S71M) to cytoplasmic actin measured by fluorescence anisotropy competition assays. Fluorescence anisotropy of Atto488-WAVE1_WCA_ [4 nM] as a function of increasing profilin1 (wt, S71M as indicated) concentration in the presence of a constant amount of actin monomers [150 nM]. Lines are fits to an exact analytical competition model (see Methods). Each data point represents the mean value from three independent experiments. Error indicators are SD. (F) Barbed end growth velocities measured from TIRF-M single filament assays using profilin1 (S71M):actin complexes. Points represent the calculated mean of the actin filament elongation rate at a given profilin-actin concentration [N ≥40 for each condition, error bars = SD]. Continuous lines are hyperbolic fits yielding the indicated maximum filament growth rates at saturation (See Methods). Inset: Kymographs of representative individual filaments at 150 μM profilin-actin.

**Fig. S4:**
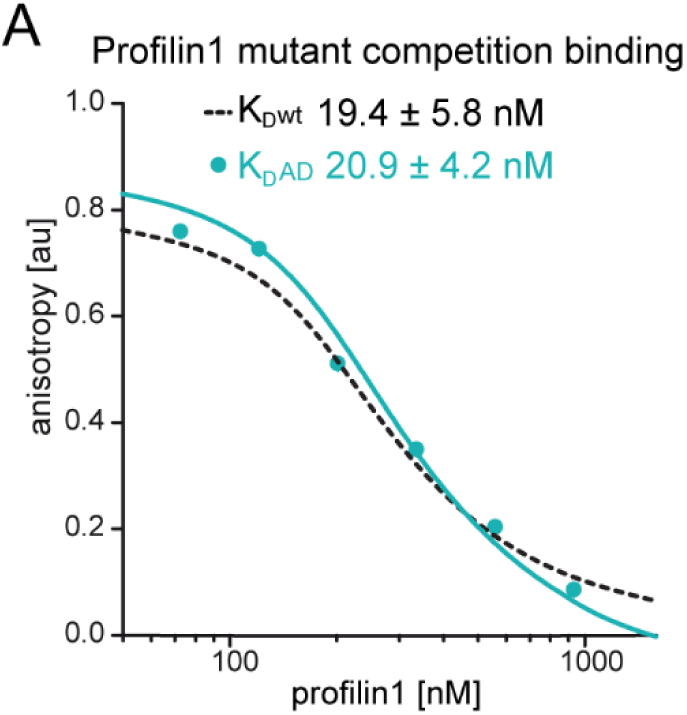
Affinity measurements of profilin1 to wt β-actin and ATPase-deficient actin. (A) Binding of profilin1 to cytoplasmic actin, either wt (black dashed) or AD (cyan) measured by fluorescence anisotropy competition assays. Fluorescence anisotropy of Atto488-WAVE1_WCA_ [4 nM] as a function of increasing profilin1 concentration in the presence of a constant amount of actin monomers [150 nM]. Lines are fits to an exact analytical competition model (see Methods). Each data point represents the mean value from three independent experiments. Error indicators are SD.

**Fig. S5:**
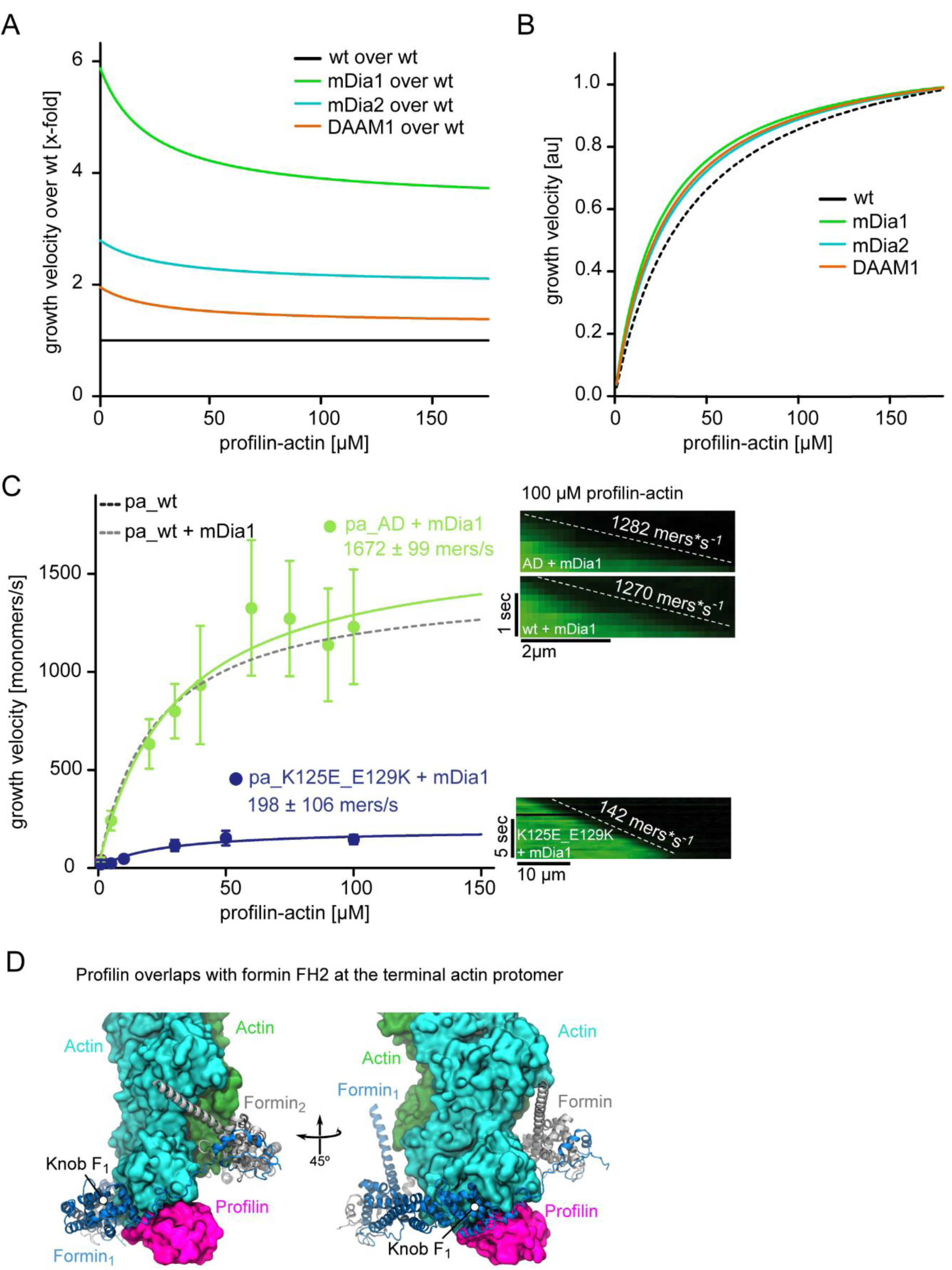
Profilin release but not ATP hydrolysis is limiting for formin-mediated actin polymerization. (A) The relative (x-fold) enhancement of the filament growth velocity by either mDia1, mDia2 or DAAM1 as indicated. Growth velocities (Fig. 5C) were normalized by the speed of actin growth in the absence of formins. (B) Growth velocities in the presence of mDia1, mDia2 or DAAM1 as indicated (Fig. 5C) normalized by their characteristic maximal growth rate. This re-normalization shows that all formins lower the concentration of profilin-actin required for half-maximal elongation speeds and thus slightly broaden the region of concentration invariance. (C) mDia1-catalyzed barbed end growth velocities of profilin1–actin (either both proteins wt (dashed), or AD actin (light green) or tight binding profilin-1(K125E+E129K) (cyan)) from TIRF-M assays. Note that the tight binding profilin-mutant reduces the maximal elongation rate in contrast to ATPase-deficient actin. Points represent the calculated mean of the actin filament elongation rate at a given profilin-actin concentration [N ≥35 for each condition, error bars = SD]. Lines are hyperbolic fits. Right: Kymographs of the growth of representative individual filaments at 100 μM profilin-actin. (D) Model of the FH2 domain of mDia1 (blue) bound to the barbed end of an actin filament bound to profilin (see Methods). The actin subunits (green) and profilin (magenta) are shown as surfaces. The knob region of the FH2 domain contacts the profilin molecule bound at the terminal protomer of the filament.

**Fig. S6:**
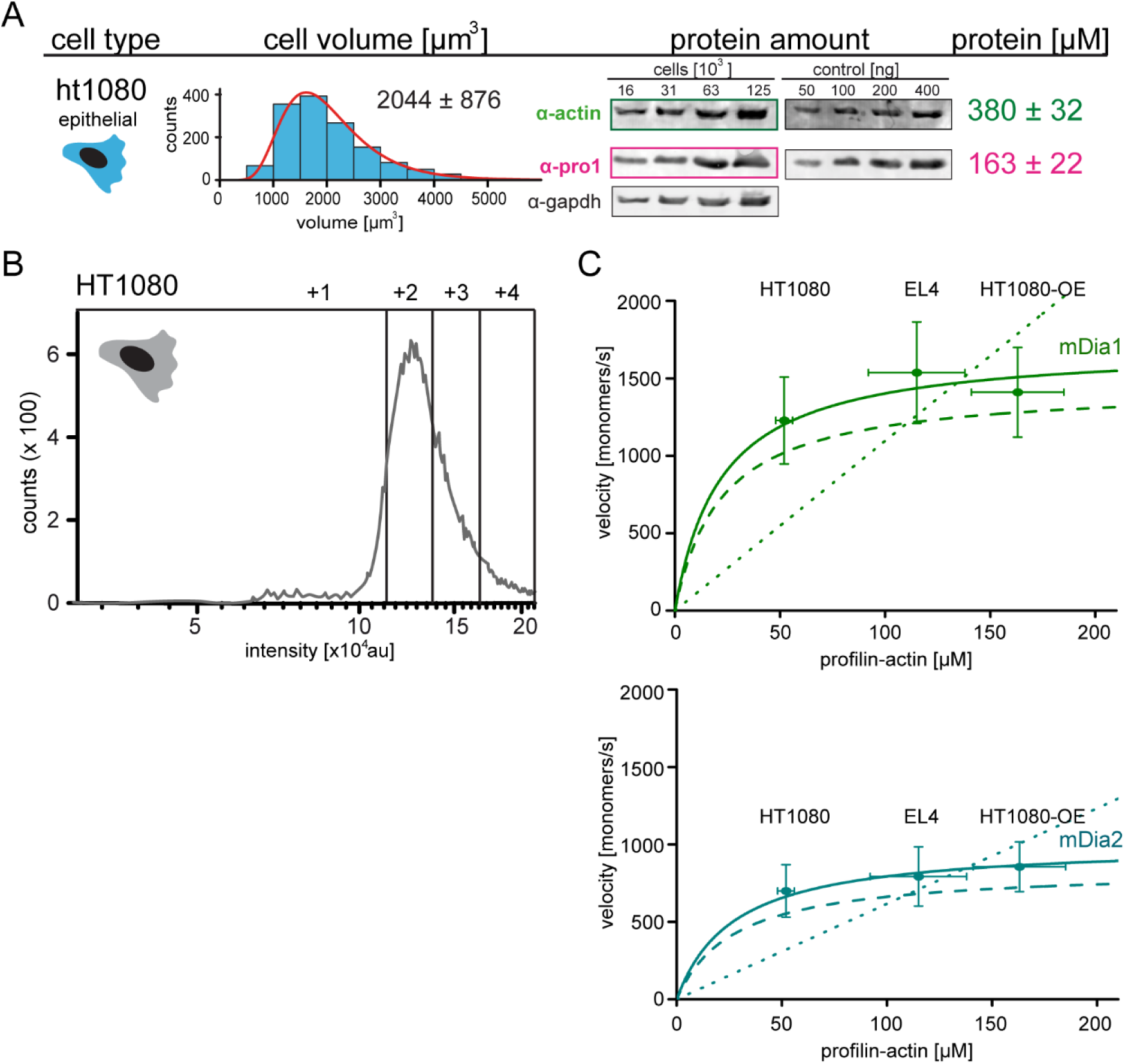
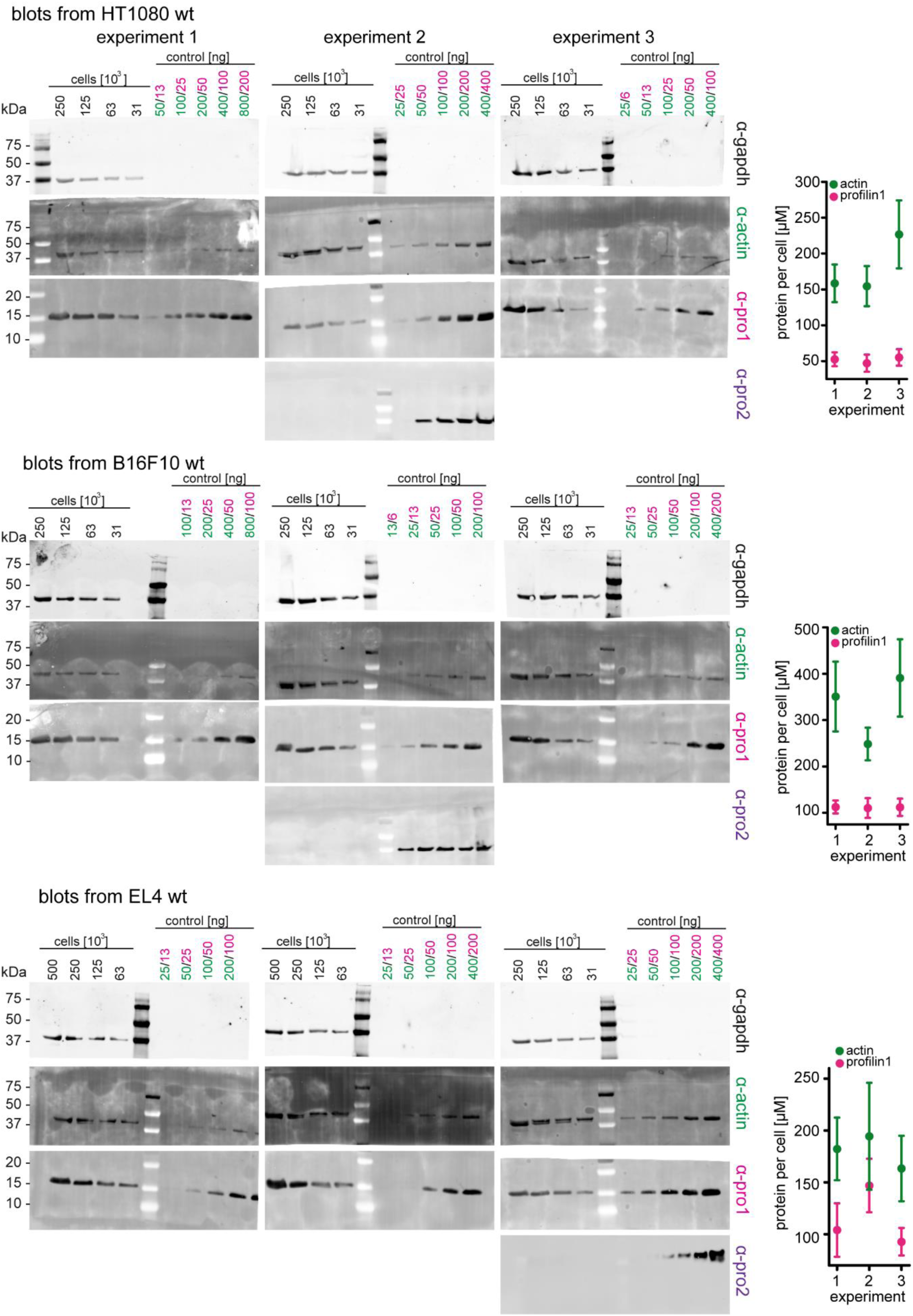
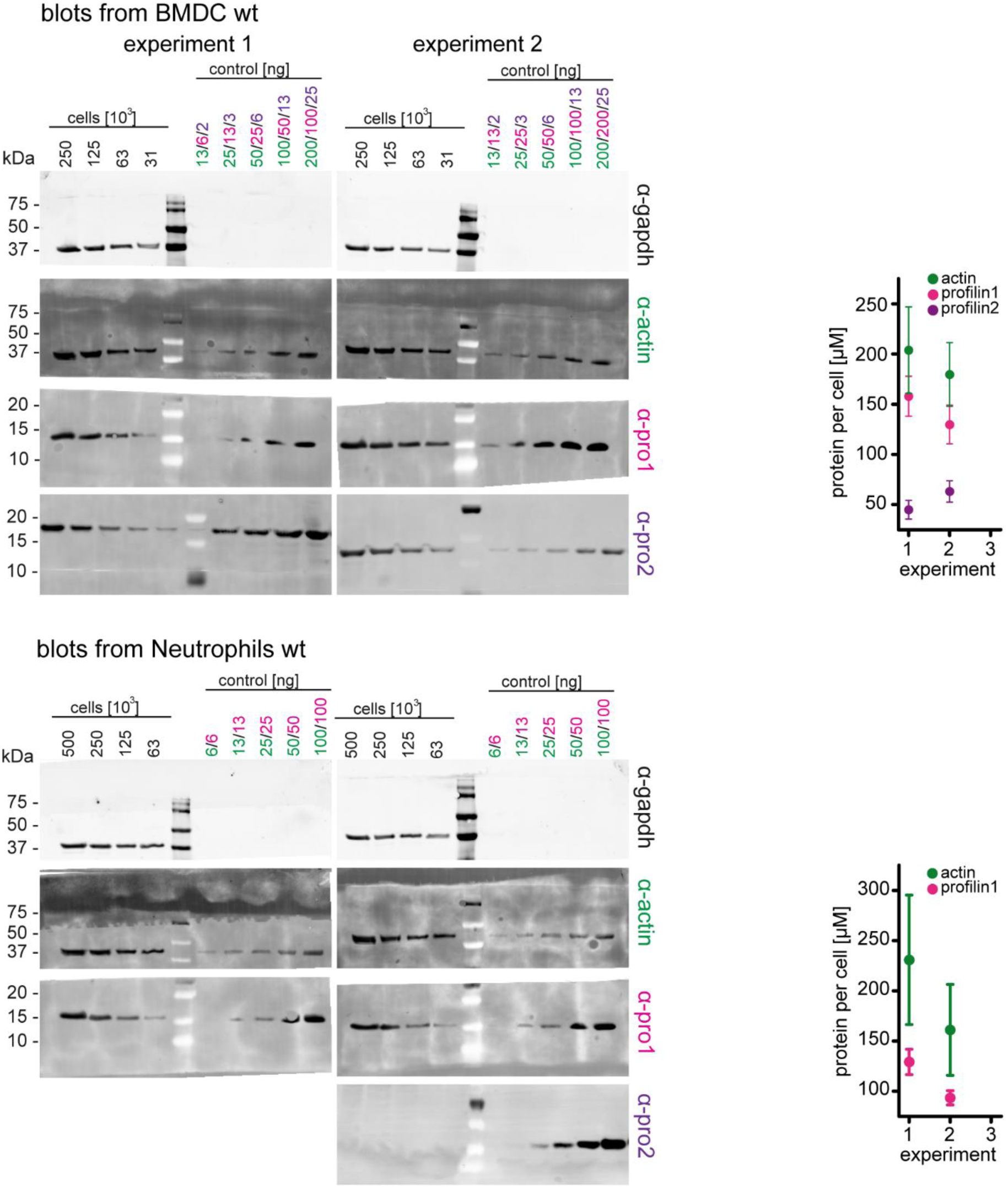
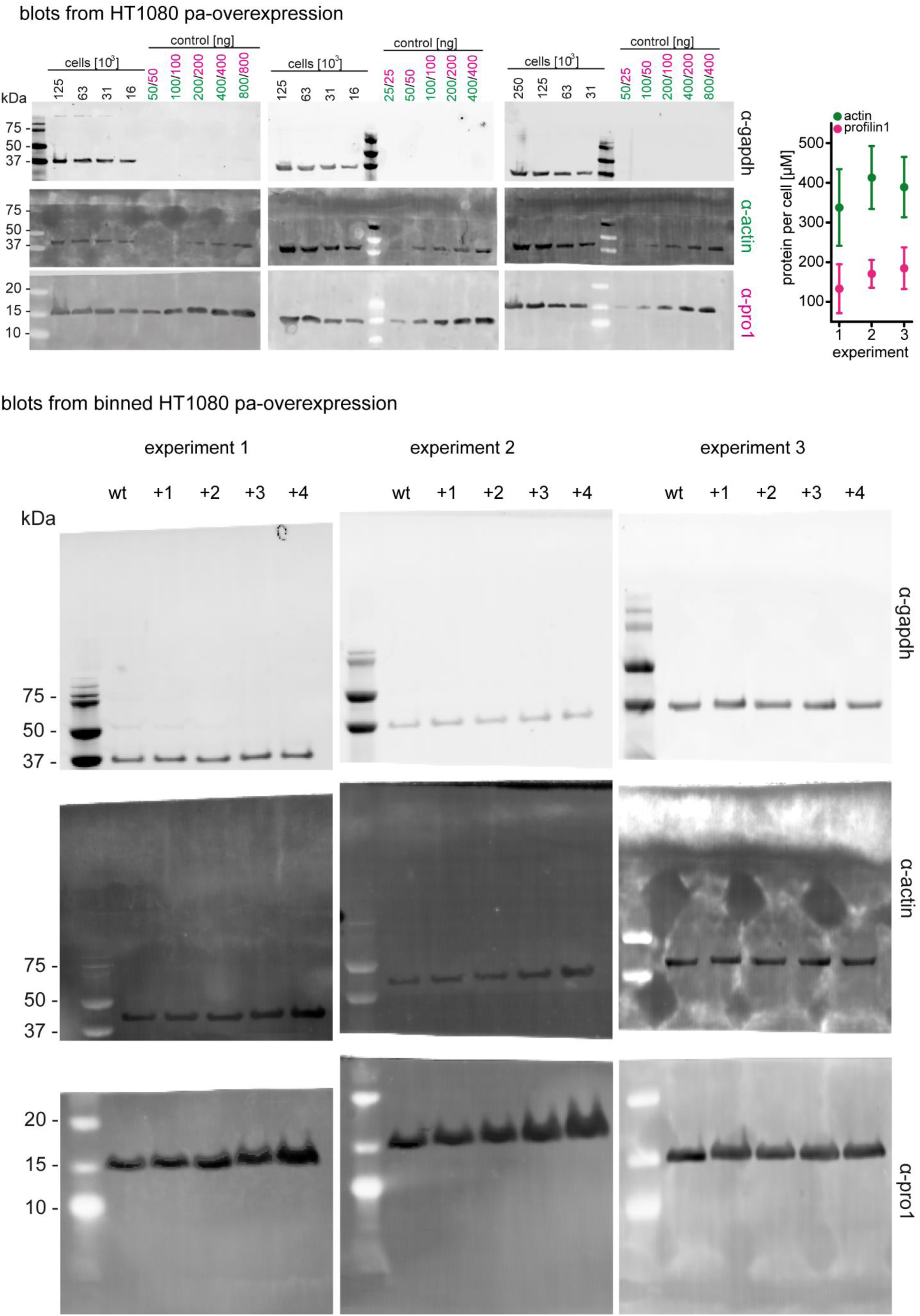
Profilin1-actin overexpression and in vivo formin speeds at different profilin1-actin levels. (A) Profilin-actin concentration determination in mammalian HT1080 cells overexpressing profilin1 and β-actin. Left to right: name of cell type and origin, histograms of the single cell volume from fluorescence eXclusion measurements, quantitative western blot analysis of cellular actin, profilin1 amount (left: titration of cell number, right: standard curve of recombinant proteins), calculated mean protein concentration [μM] per cell with experimental error (SD, N=3 independent experiments, see Methods). (B) Histogram from FACS analysis showing the ScarletI intensity distribution from a polyclonal population of HT1080 cells overexpressing profilin1, mScarletI and β-actin. This population was sorted into subpopulations as indicated (+1 to +4). (C) Mean velocities of either mDia1 (top) and mDia2 (bottom) in different mammalian cell types (wt HT080, EL4 and HT1080 overexpressing profilin-actin) plotted as a function of the quantified profilin-actin concentration. Error indicators are SD. Dashed lines are the formin speeds determined in vitro (see Fig. 5C), dotted lines are linear fits through the origin to the in vivo data. Continuous lines are fits to the in vivo data by a hyperbolic model with only one free parameter (v_max_, the maximal growth velocity). K_0.5_ (the profilin-actin concentration at half-maximal elongation speed) was fixed to the value determined in vitro (see Methods). **Western blots and graphical summary of profilin-actin levels per cell.** Determination of protein-levels in different cell types by quantitative western blot analysis. For each cell type (HT1080, B16F10, BMDC, neutrophils, HT1080/B16F10 overexpressing profilin1 and β-actin) the amount of total profilin1/2 and actin was detected by monoclonal antibodies and calculated with in vitro protein controls in multiple experiments (see Methods). The amount of profilin2 control is equal to the amount of profilin1 unless otherwise indicated. Right: Calculated profilin1/2 and actin concentrations per cell for individual experiments, errors indicate the SD. Last: Western blots from wt HT1080 cells and HT1080 overexpressing profilin1 and β-actin from a polyclonal integrated construct (+1 to +4) after sorting them by their mScarletI fluorescence intensity (see also Fig. S6B). For each sample, 250.000 cells were applied.

## Supplementary Movie Legends

Movie S1

Polymerization of actin filaments from different profilin1-actin concentrations. Filaments were visualized with 10 nM Cy5-UTRN_261_ in TIRF-M. Polymerization from increasing profilin1-actin concentrations from left to right: 2.5 μM, 40 μM, 125 μM, 175 μM.

Movie S2

Polymerization of actin filaments from different profilin1 mutant-actin complexes at 125 μM. Filaments were visualized with 10 nM Cy5-UTRN_261_ in TIRF-M. Polymerization was performed from the following profilin1 mutant-actin complexes, left to right: profilin1-K125E + E129K, -wt, -E82A, -R88K.

Movie S3

Polymerization of actin filaments from profilin1-actin at different concentrations in presence of mDia1 FH1-FH2. Filaments were acquired in TIRF-M (filaments with 10 nM Cy5-UTRN_261_ - green; 0.7 nM TMR-mDia1 FH-FH2 – magenta). mDia1 mediated actin filament barbed end polymerization was performed at different profilin1-actin concentrations, left to right: 1 μM, 10 μM, 20 μM. For guidance, an example of a visible labeled mDia1 molecule processively moving with a filament barbed end is highlighted with an error.

Movie S4

Polymerization of actin filaments from 75 μM profilin1-actin in presence/absence of formins. Filaments were visualized with 10 nM Cy5-UTRN_261_ in TIRF-M. All filament barbed ends were saturated with 15 nM formin FH1-FH2. Polymerization was performed in presence of different formins, left to right: wt (no formin), + DAAM1, +mDia2, +mDia1.

Movie S5

mDia1 and mDia2 formin single molecule movement in HT1080 cells under conditions with either wt or overexpression of profilin1–actin. mNeonGreen–mDia1/2 FH1-FH2 single molecules were visualized in TIRF-M. To indicate the cell shape, HT1080 cells were masked. Top: mDia1 (left) and mDia2 (right) molecules in wt HT1080 cells. Bottom: mDia1 and mDia2 molecules in HT1080 cells overexpressing profilin and actin.

Movie S6

In vivo mDia2 single molecule movement in absence/presence of latrunculinB, JASP and y27632. To indicate the cell shape, HT1080 cells were masked. mNeonGreen-mDia2 FH1-FH2 single molecules were visualized in TIRF-M. mDia2 molecules were monitored without and after 10 min of drug treatment. The following drugs were applied to the cells, left to right: no drug treatment, 500 nM latrunculinB (latB), 8 μM JASP, 10 μM y27632.

## References

Aydin F, Courtemanche N, Pollard TD, Voth GA. 2018. Gating mechanisms during actin filament elongation by formins. Elife 7. doi:10.7554/eLife.37342

Bieling P, Hansen SD, Akin O, Li T-D, Hayden CC, Fletcher DA, Mullins RD. 2018. WH2 and proline-rich domains of WASP-family proteins collaborate to accelerate actin filament elongation. EMBO J 37:102–121. doi:10.15252/embj.201797039

Bieling P, Li T-D, Weichsel J, McGorty R, Jreij P, Huang B, Fletcher DA, Mullins RD. 2016. Force Feedback Controls Motor Activity and Mechanical Properties of Self-Assembling Branched Actin Networks. Cell 164:115–127. doi:10.1016/j.cell.2015.11.057

Blanchoin L, Boujemaa-Paterski R, Sykes C, Plastino J. 2014. Actin dynamics, architecture, and mechanics in cell motility. Physiol Rev 94:235–263. doi:10.1152/physrev.00018.2013

Blanchoin L, Pollard TD. 2002. Hydrolysis of ATP by polymerized actin depends on the bound divalent cation but not profilin. Biochemistry 41:597–602.

Cadart C, Zlotek-Zlotkiewicz E, Venkova L, Thouvenin O, Racine V, Le Berre M, Monnier S, Piel M. 2017. Fluorescence eXclusion Measurement of volume in live cells. Methods Cell Biol 139:103–120. doi:10.1016/bs.mcb.2016.11.009

Carlier M-F, Shekhar S. 2017. Global treadmilling coordinates actin turnover and controls the size of actin networks. Nat Rev Mol Cell Biol 18:389–401. doi:10.1038/nrm.2016.172

Courtemanche N. 2018. Mechanisms of formin-mediated actin assembly and dynamics. Biophys Rev 10:1553–1569. doi:10.1007/s12551-018-0468-6

Courtemanche N, Pollard TD. 2013. Interaction of profilin with the barbed end of actin filaments. Biochemistry 52:6456–6466. doi:10.1021/bi400682n

Dimchev G, Steffen A, Kage F, Dimchev V, Pernier J, Carlier M-F, Rottner K. 2017. Efficiency of lamellipodia protrusion is determined by the extent of cytosolic actin assembly. Mol Biol Cell 28:1311–1325. doi:10.1091/mbc.E16-05-0334

Gutsche-Perelroizen I, Lepault J, Ott A, Carlier MF. 1999. Filament assembly from profilin-actin. J Biol Chem 274:6234–6243.

Hatano T, Alioto S, Roscioli E, Palani S, Clarke ST, Kamnev A, Hernandez-Fernaud JR, Sivashanmugam L, Chapa-Y-Lazo B, Jones AME, Robinson RC, Sampath K, Mishima M, McAinsh AD, Goode BL, Balasubramanian MK. 2018. Rapid production of pure recombinant actin isoforms in Pichia pastoris. J Cell Sci 131. doi:10.1242/jcs.213827

Higashida C, Kiuchi T, Akiba Y, Mizuno H, Maruoka M, Narumiya S, Mizuno K, Watanabe N. 2013. F- and G-actin homeostasis regulates mechanosensitive actin nucleation by formins. Nat Cell Biol 15:395–405. doi:10.1038/ncb2693

Higashida C, Miyoshi T, Fujita A, Oceguera-Yanez F, Monypenny J, Andou Y, Narumiya S, Watanabe N. 2004. Actin polymerization-driven molecular movement of mDia1 in living cells. Science 303:2007–2010. doi:10.1126/science.1093923

Jégou A, Carlier M-F, Romet-Lemonne G. 2013. Formin mDia1 senses and generates mechanical forces on actin filaments. Nat Commun 4:1883. doi:10.1038/ncomms2888

Jégou A, Niedermayer T, Orbán J, Didry D, Lipowsky R, Carlier M-F, Romet-Lemonne G. 2011. Individual actin filaments in a microfluidic flow reveal the mechanism of ATP hydrolysis and give insight into the properties of profilin. PLoS Biol 9:e1001161. doi:10.1371/journal.pbio.1001161

Kaiser DA, Vinson VK, Murphy DB, Pollard TD. 1999. Profilin is predominantly associated with monomeric actin in Acanthamoeba. J Cell Sci 112 (**Pt 21**):3779–3790.

Kinosian HJ, Selden LA, Gershman LC, Estes JE. 2002. Actin filament barbed end elongation with nonmuscle MgATP-actin and MgADP-actin in the presence of profilin. Biochemistry 41:6734– 6743.

Kinosian HJ, Selden LA, Gershman LC, Estes JE. 2000. Interdependence of profilin, cation, and nucleotide binding to vertebrate non-muscle actin. Biochemistry 39:13176–13188.

Koestler SA, Rottner K, Lai F, Block J, Vinzenz M, Small JV. 2009. F- and G-actin concentrations in lamellipodia of moving cells. PLoS ONE 4:e4810. doi:10.1371/journal.pone.0004810

Kovar DR, Harris ES, Mahaffy R, Higgs HN, Pollard TD. 2006. Control of the assembly of ATP- and ADP-actin by formins and profilin. Cell 124:423–435. doi:10.1016/j.cell.2005.11.038

Merino F, Pospich S, Funk J, Wagner T, Küllmer F, Arndt H-D, Bieling P, Raunser S. 2018. Structural transitions of F-actin upon ATP hydrolysis at near-atomic resolution revealed by cryo-EM. Nat Struct Mol Biol 25:528–537. doi:10.1038/s41594-018-0074-0

Mogilner A, Oster G. 1996. Cell motility driven by actin polymerization. Biophys J 71:3030–3045. doi:10.1016/S0006-3495(96)79496-1

Mouneimne G, Hansen SD, Selfors LM, Petrak L, Hickey MM, Gallegos LL, Simpson KJ, Lim J, Gertler FB, Hartwig JH, Mullins RD, Brugge JS. 2012. Differential remodeling of actin cytoskeleton architecture by profilin isoforms leads to distinct effects on cell migration and invasion. Cancer Cell 22:615–630. doi:10.1016/j.ccr.2012.09.027

Mueller J, Szep G, Nemethova M, de Vries I, Lieber AD, Winkler C, Kruse K, Small JV, Schmeiser C, Keren K, Hauschild R, Sixt M. 2017. Load Adaptation of Lamellipodial Actin Networks. Cell 171:188–200.e16. doi:10.1016/j.cell.2017.07.051

Ohki T, Ohno C, Oyama K, Mikhailenko SV, Ishiwata S. 2009. Purification of cytoplasmic actin by affinity chromatography using the C-terminal half of gelsolin. Biochem Biophys Res Commun 383:146–150. doi:10.1016/j.bbrc.2009.03.144

Oosawa F, Asakura S. 1975. Thermodynamics of the Polymerization of Protein, Molecular biology. Academic Press.

Pantaloni D, Carlier MF. 1993. How profilin promotes actin filament assembly in the presence of thymosin beta 4. Cell 75:1007–1014.

Paul AS, Pollard TD. 2009. Review of the mechanism of processive actin filament elongation by formins. Cell Motil Cytoskeleton 66:606–617. doi:10.1002/cm.20379

Pernier J, Shekhar S, Jegou A, Guichard B, Carlier M-F. 2016. Profilin Interaction with Actin Filament Barbed End Controls Dynamic Instability, Capping, Branching, and Motility. Dev Cell 36:201– 214. doi:10.1016/j.devcel.2015.12.024

Pollard TD. 1986. Rate constants for the reactions of ATP- and ADP-actin with the ends of actin filaments. J Cell Biol 103:2747–2754.

Pollard TD, Blanchoin L, Mullins RD. 2000. Molecular mechanisms controlling actin filament dynamics in nonmuscle cells. Annu Rev Biophys Biomol Struct 29:545–576. doi:10.1146/annurev.biophys.29.1.545

Pollard TD, Cooper JA. 1984. Quantitative analysis of the effect of Acanthamoeba profilin on actin filament nucleation and elongation. Biochemistry 23:6631–6641.

Pring M, Weber A, Bubb MR. 1992. Profilin-actin complexes directly elongate actin filaments at the barbed end. Biochemistry 31:1827–1836.

Raz-Ben Aroush D, Ofer N, Abu-Shah E, Allard J, Krichevsky O, Mogilner A, Keren K. 2017. Actin Turnover in Lamellipodial Fragments. Curr Biol 27:2963–2973.e14. doi:10.1016/j.cub.2017.08.066

Renkawitz J, Schumann K, Weber M, Lämmermann T, Pflicke H, Piel M, Polleux J, Spatz JP, Sixt M. 2009. Adaptive force transmission in amoeboid cell migration. Nat Cell Biol 11:1438–1443. doi:10.1038/ncb1992

Romero S, Didry D, Larquet E, Boisset N, Pantaloni D, Carlier M-F. 2007. How ATP hydrolysis controls filament assembly from profilin-actin: implication for formin processivity. J Biol Chem 282:8435–8445. doi:10.1074/jbc.M609886200

Romero S, Le Clainche C, Didry D, Egile C, Pantaloni D, Carlier M-F. 2004. Formin is a processive motor that requires profilin to accelerate actin assembly and associated ATP hydrolysis. Cell 119:419–429. doi:10.1016/j.cell.2004.09.039

Rotty JD, Wu C, Haynes EM, Suarez C, Winkelman JD, Johnson HE, Haugh JM, Kovar DR, Bear JE. 2015. Profilin-1 serves as a gatekeeper for actin assembly by Arp2/3-dependent and -independent pathways. Dev Cell 32:54–67. doi:10.1016/j.devcel.2014.10.026

Schutt CE, Myslik JC, Rozycki MD, Goonesekere NC, Lindberg U. 1993. The structure of crystalline profilin-beta-actin. Nature 365:810–816. doi:10.1038/365810a0

Skruber K, Read T-A, Vitriol EA. 2018. Reconsidering an active role for G-actin in cytoskeletal regulation. J Cell Sci 131. doi:10.1242/jcs.203760

Suarez C, Carroll RT, Burke TA, Christensen JR, Bestul AJ, Sees JA, James ML, Sirotkin V, Kovar DR. 2015. Profilin regulates F-actin network homeostasis by favoring formin over Arp2/3 complex. Dev Cell 32:43–53. doi:10.1016/j.devcel.2014.10.027

Suarez C, Kovar DR. 2016. Internetwork competition for monomers governs actin cytoskeleton organization. Nat Rev Mol Cell Biol 17:799–810. doi:10.1038/nrm.2016.106

Vavylonis D, Kovar DR, O’Shaughnessy B, Pollard TD. 2006. Model of formin-associated actin filament elongation. Mol Cell 21:455–466. doi:10.1016/j.molcel.2006.01.016

Vinson VK, De La Cruz EM, Higgs HN, Pollard TD. 1998. Interactions of Acanthamoeba profilin with actin and nucleotides bound to actin. Biochemistry 37:10871–10880. doi:10.1021/bi980093l

Witke W, Sutherland JD, Sharpe A, Arai M, Kwiatkowski DJ. 2001. Profilin I is essential for cell survival and cell division in early mouse development. Proc Natl Acad Sci USA 98:3832–3836. doi:10.1073/pnas.051515498

## Supplemental References

Aitken CE, Marshall RA, Puglisi JD. 2008. An oxygen scavenging system for improvement of dye stability in single-molecule fluorescence experiments. Biophys J 94:1826–1835. doi:10.1529/biophysj.107.117689

Berger S, Procko E, Margineantu D, Lee EF, Shen BW, Zelter A, Silva D-A, Chawla K, Herold MJ, Garnier J-M, Johnson R, MacCoss MJ, Lessene G, Davis TN, Stayton PS, Stoddard BL, Fairlie WD, Hockenbery DM, Baker D. 2016. Computationally designed high specificity inhibitors delineate the roles of BCL2 family proteins in cancer. Elife 5. doi:10.7554/eLife.20352

Bieling P, Telley IA, Hentrich C, Piehler J, Surrey T. 2010. Fluorescence microscopy assays on chemically functionalized surfaces for quantitative imaging of microtubule, motor, and +TIP dynamics. Methods Cell Biol 95:555–580. doi:10.1016/S0091-679X(10)95028-0

Crooks GE, Hon G, Chandonia J-M, Brenner SE. 2004. WebLogo: a sequence logo generator. Genome Res 14:1188–1190. doi:10.1101/gr.849004

Duellberg C, Cade NI, Holmes D, Surrey T. 2016. The size of the EB cap determines instantaneous microtubule stability. Elife 5. doi:10.7554/eLife.13470

Edelstein AD, Tsuchida MA, Amodaj N, Pinkard H, Vale RD, Stuurman N. 2014. Advanced methods of microscope control using μManager software. J Biol Methods 1. doi:10.14440/jbm.2014.36

Fleishman SJ, Leaver-Fay A, Corn JE, Strauch E-M, Khare SD, Koga N, Ashworth J, Murphy P, Richter F, Lemmon G, Meiler J, Baker D. 2011. RosettaScripts: a scripting language interface to the Rosetta macromolecular modeling suite. PLoS ONE 6:e20161. doi:10.1371/journal.pone.0020161

Hudson EN, Weber G. 1973. Synthesis and characterization of two fluorescent sulfhydryl reagents. Biochemistry 12:4154–4161.

Leaver-Fay A, Tyka M, Lewis SM, Lange OF, Thompson J, Jacak R, Kaufman K, Renfrew PD, Smith CA, Sheffler W, Davis IW, Cooper S, Treuille A, Mandell DJ, Richter F, Ban Y-EA, Fleishman SJ, Corn JE, Kim DE, Lyskov S, Berrondo M, Mentzer S, Popović Z, Havranek JJ, Karanicolas J, Das R, Meiler J, Kortemme T, Gray JJ, Kuhlman B, Baker D, Bradley P. 2011. ROSETTA3: an object-oriented software suite for the simulation and design of macromolecules. Meth Enzymol 487:545–574. doi:10.1016/B978-0-12-381270-4.00019-6

Miki M, Barden JA, dos Remedios CG, Phillips L, Hambly BD. 1987. Interaction of phalloidin with chemically modified actin. Eur J Biochem 165:125–130.

Nezami A, Poy F, Toms A, Zheng W, Eck MJ. 2010. Crystal structure of a complex between amino and carboxy terminal fragments of mDia1: insights into autoinhibition of diaphanous-related formins. PLoS ONE 5. doi:10.1371/journal.pone.0012992

Otomo T, Tomchick DR, Otomo C, Panchal SC, Machius M, Rosen MK. 2005. Structural basis of actin filament nucleation and processive capping by a formin homology 2 domain. Nature 433:488–494. doi:10.1038/nature03251

Peng GE, Wilson SR, Weiner OD. 2011. A pharmacological cocktail for arresting actin dynamics in living cells. Mol Biol Cell 22:3986–3994. doi:10.1091/mbc.E11-04-0379

Pollard TD. 1982. Myosin purification and characterization. Methods Cell Biol 24:333–371.

Rasnik I, McKinney SA, Ha T. 2006. Nonblinking and long-lasting single-molecule fluorescence imaging. Nat Methods 3:891–893. doi:10.1038/nmeth934

Shimada A, Nyitrai M, Vetter IR, Kühlmann D, Bugyi B, Narumiya S, Geeves MA, Wittinghofer A. 2004. The core FH2 domain of diaphanous-related formins is an elongated actin binding protein that inhibits polymerization. Mol Cell 13:511–522.

Tinevez J-Y, Perry N, Schindelin J, Hoopes GM, Reynolds GD, Laplantine E, Bednarek SY, Shorte SL, Eliceiri KW. 2017. TrackMate: An open and extensible platform for single-particle tracking. Methods 115:80–90. doi:10.1016/j.ymeth.2016.09.016

Vargas P, Maiuri P, Bretou M, Sáez PJ, Pierobon P, Maurin M, Chabaud M, Lankar D, Obino D, Terriac E, Raab M, Thiam H-R, Brocker T, Kitchen-Goosen SM, Alberts AS, Sunareni P, Xia S, Li R, Voituriez R, Piel M, Lennon-Duménil A-M. 2016. Corrigendum: Innate control of actin nucleation determines two distinct migration behaviours in dendritic cells. Nat Cell Biol 18:234. doi:10.1038/ncb3301

Wang ZX. 1995. An exact mathematical expression for describing competitive binding of two different ligands to a protein molecule. FEBS Lett 360:111–114.

Webb B, Sali A. 2016. Comparative Protein Structure Modeling Using MODELLER. Curr Protoc Bioinformatics 54:5.6.1-5.6.37. doi:10.1002/cpbi.3

Zalevsky J, Grigorova I, Mullins RD. 2001. Activation of the Arp2/3 complex by the Listeria acta protein. Acta binds two actin monomers and three subunits of the Arp2/3 complex. J Biol Chem 276:3468–3475. doi:10.1074/jbc.M006407200

